# STAT3 dual role in sex-biased and differentiation-gated interferon gamma memory

**DOI:** 10.64898/2026.10.21.760183

**Authors:** Justyna Topa, Wojciech Snoch, Mateusz Modzelewski, Sohaib Khan, Elzbieta Chrusciel, Artur Pirog, Sachin Kote, Wojciech Siwek

**Affiliations:** International Centre for Cancer Vaccine Science, University of Gdansk, PL; Department of Molecular Biology, Massachusetts General Hospital Research Institute & Department of Genetics, Harvard Medical School, Boston, USA

**Keywords:** Interferon-Gamma Memory, STAT Transcription Factors, Sex-Biased Immune Response, Transcriptional Precision, Monocyte-to-Macrophage Differentiation, Trained Immunity, Epigenetics

## Abstract

The innate immune system possesses a unique form of memory, known as trained immunity. An initial stimulus induces cellular reprogramming for a non-specific amplified response to a subsequent challenge. This process can be achieved through gene priming, where exposure to a specific cue sensitizes future gene expression. Interferon-gamma (IFNγ) memory is a specialized form of priming, where both the initial and recall signals are mediated by the same IFNγ cytokine. The mechanisms governing IFNγ memory in humans remain poorly understood. We investigated the mechanisms governing IFNγ memory and characterized it in human primary macrophages. We discovered a dual role for the STAT3 transcription factor in IFNγ memory. It acts as a repressor of memory establishment by indirectly inhibiting STAT1 Ser727 phosphorylation, while also being required for memory recall. Furthermore, we demonstrated that IFNγ memory is sex-biased, with males exhibiting a more pronounced effect characterized by reduced variability in gene expression upon re-exposure to IFNγ. This defines IFNγ recall as a function of transcriptional precision. We also identified a novel family of chemokine genes that exhibit IFNγ concentration dependent memory effects. Additionally, we established a model system to study this process and found that memory is critically dependent on the cellular phenotype. Specifically, our results show that for some genes, monocyte-to-macrophage differentiation is necessary, while for others, this differentiation step is sufficient for recall of the previous response.

Our findings redefine the IFNγ memory program, establishing it as a sex-biased phenomenon governed by a novel regulatory network involving differentiation and STAT3 function. These insights provide critical context for understanding immune heterogeneity and are essential for the rational design of personalized therapies that harness trained immunity.

**Graphical Abstract:** 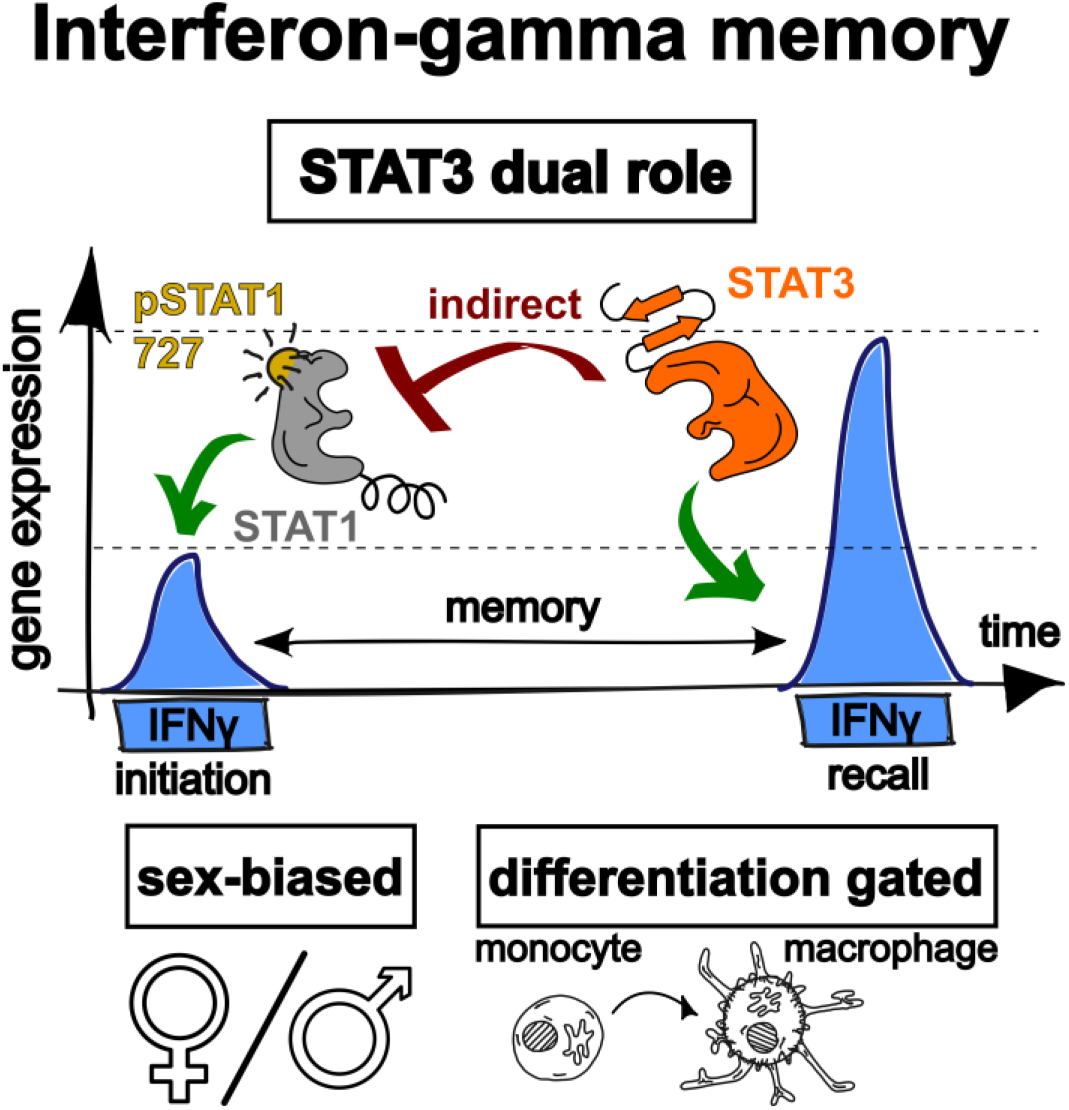

**Highlights:**

- STAT3 indirectly inhibits IFNγ memory initiation by repressing STAT1 Ser727 phosphorylation, but is necessary for memory recall.
- IFNγ memory exhibits a sex bias, with males demonstrating a more pronounced effect, characterized by reduced variability in gene expression upon re-exposure to IFNγ.
- Monocyte-to-macrophage differentiation controls recall of IFNγ memory, suggesting a critical role of cellular maturation in memory regulation.

## Introduction

The immune system ability to remember past encounters with pathogens is a crucial aspect of host defense. Trained immunity^1^ or innate immune memory^2^ enables innate immune cells to acquire long-term functional adaptations after exposure to pathogens or danger signals. This phenomenon enhances responsiveness upon rechallenge and has profound implications for host defense against recurrent infections, vaccine efficacy and the pathogenesis of chronic inflammatory diseases^3^. Unlike adaptive immunity, trained immunity does not rely on antigen-specific receptors but instead involves epigenetic and transcriptional reprogramming^4^. Beyond trained immunity, the innate immune system can also dampen its response through a process called tolerance^5^. This occurs when repeated exposure to specific molecules leads to decreased or abolished responses, thereby preventing excessive or unwarranted inflammation and promoting host homeostasis^6^. Understanding the molecular underpinnings of these processes is critical for developing novel immunotherapies and improving clinical outcomes in infections, autoimmune disorders, and oncology.

Trained immunity often operates via gene priming, where specific genes are pre-activated or poised for a changed transcriptional response upon re-exposure to stimuli, often with a different signal^7,8^. Epigenetic memory is a special form of priming in which genes show increased rates of expression after re-stimulation^9^. This phenomenon has been identified in various contexts, including cytokine-induced pathways like interferon gamma (IFNγ) signaling^10,11^. In our previous work, we focused on epigenetic memory where the initial and following stimulations are the same IFNγ signal. We used Guanylate Binding Proteins 5 (GBP5) and 1 (GBP1), activators of inflammation^12,13^, as our key model memory genes (Figure 1A)^14,15^.

**Figure 1.**
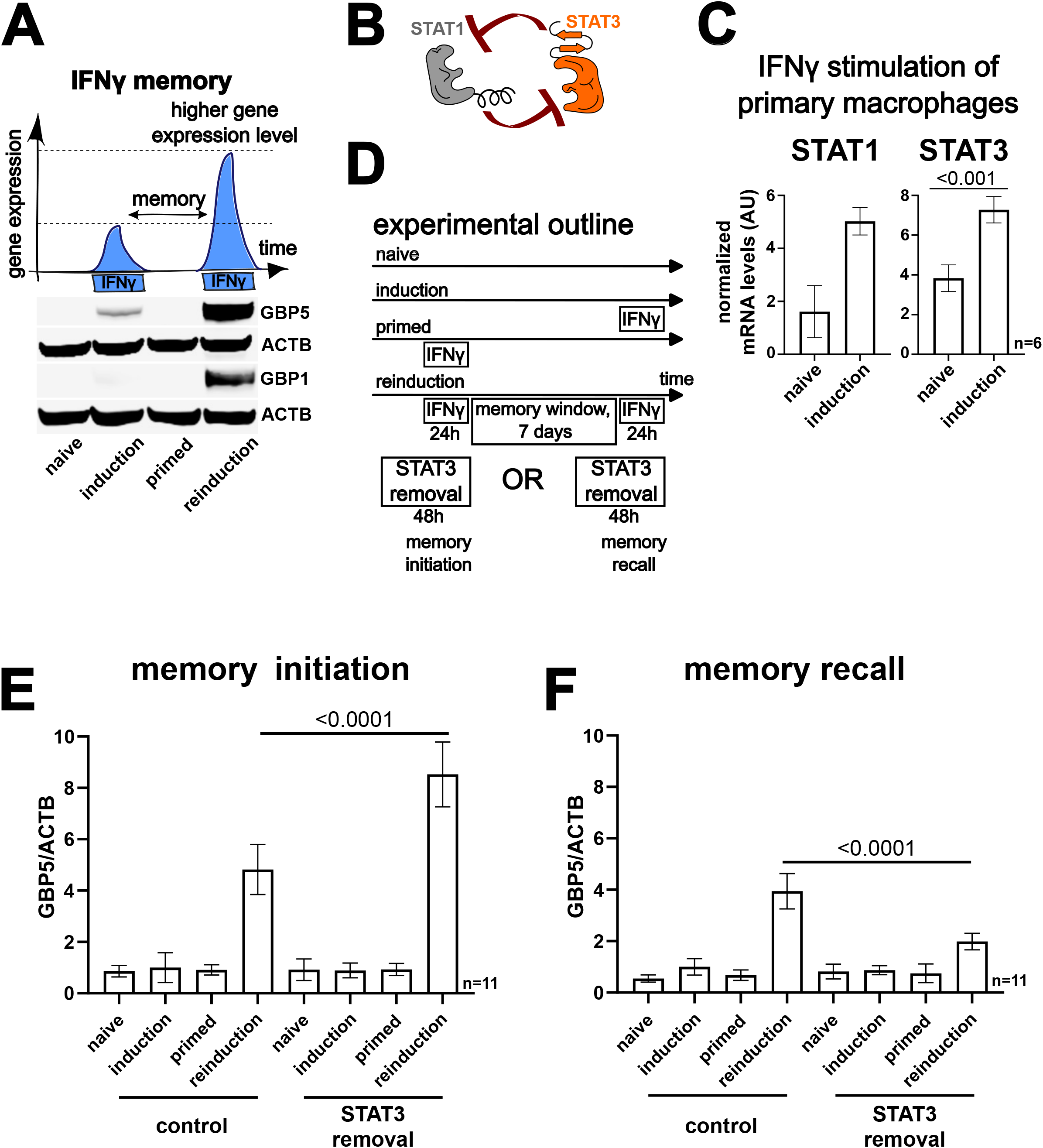
STAT3 is an inhibitor of interferon gamma (IFNγ) memory, but is required for recall. **(A)** Overview of IFNγ memory (top), western blot results showing the effect in HeLa cells for key memory genes: GBP5 and GBP1 (bottom). ACTB – loading control. **(B)** Graphical representation of cross-inhibitory functions of STAT1 and STAT3. **(C)** Expression level of STAT1 and STAT3 in naïve and IFNγ stimulated (50 ng/mL) human primary macrophages. **(D)** Experimental scheme to investigate the effect of transient removal of STAT3 using 10µM SD36 PROTAC on the initiation and recall of IFNγ memory in HeLa cells. SD36 was added either before and during initial induction or before and during reinduction. **(E)** Quantification of GBP5 protein levels (normalized to ACTB; related to panel D) showing the effect of STAT3 removal on initiation and **(F)** recall of IFNγ memory.

Proposed mechanisms of IFNγ memory range from nuclear pore involvement^11^ to the cytokine sequestration at the cell membrane^16^. The current paradigm of IFNγ memory emphasizes epigenetic modifications, such as histone methylation and acetylation^17,18^, as a part of latent enhancers^7^, and DNA demethylation^19^ as key drivers of enhanced gene expression upon re-exposure. Here, we challenge this assumption by investigating how transcription factor networks regulate IFNγ memory. Interestingly, specific transcription factors were shown to maintain the memory phenotype in yeast^20,21^ and plant^22,23^ models of epigenetic memory. However, these mechanisms are poorly understood in the mammalian system. STAT family transcription factors such as STAT1 and STAT3 (Signal Transducer and Activator of Transcription 1 and 3) mediate cytokine signaling, but their roles in sustaining or modulating memory states remain largely unexplored^15^.

Canonical IFNγ signaling relies on dual phosphorylation of STAT1, Tyr701 modification is essential for homodimerization and nuclear translocation^24^, whereas phosphorylation at Ser727 is required for maximal transcriptional transactivation^25^. While STAT1 drives acute IFNγ responses, STAT3 has been implicated in both promoting and suppressing inflammatory reactions depending on context^26^. Consequently, the role of coordinated regulation of STAT1 phosphorylation dynamics and STAT3, as well as the impact of individual variability and cellular differentiation on IFNγ memory initiation and recall, requires deeper investigation.

In this study, we characterized key aspects of IFNγ memory. We found that STAT3, a transcription factor known to mediate anti-inflammatory responses^27^, plays a dual role in regulating IFNγ memory. On one hand, STAT3 inhibits memory initiation by suppressing STAT1 phosphorylation at Ser727. On the other hand, it is essential for memory recall. Moreover, by using primary human macrophages, we discovered sex-variability and stabilization-based regulation in the process. Additionally, we identified novel genes exhibiting memory and tolerance (e.g. CXCL9) as well as established a model system, using THP-1 cells, to discover the role of cellular differentiation in gating memory recall.

Together, our findings suggest that innate immune memory regulation is more dynamic and context-dependent than previously thought, involving a complex interplay between transcription factor networks and cellular differentiation. Our study provides a new framework for understanding the mechanisms governing innate immune memory and opens up avenues for development of novel immunotherapies.

## Results

### STAT3 plays a dual role in regulating IFNγ memory

Our findings indicate that STAT3 has a dual function in IFNγ memory, acting as an inhibitor of memory initiation while being necessary for recall.

We hypothesized that the STAT1-STAT3 transcription factor network, which involves mutual inhibition (Figure 1B)^26^, regulates IFNγ memory. Analysis of mRNA-seq data during IFNγ stimulation of primary macrophages showed that both STAT1 and STAT3 are induced at the mRNA level (Figure 1C). We also observed this for our experimental model, the HeLa cell line (Supplementary Figure 1A)^14^.

To understand STAT3 function in IFNγ memory, we established a quantitative dot blot assay to measure GBP5 protein levels in the HeLa cell line (Supplementary Figure 1B). We designed an experiment removing STAT3 during memory establishment or recall (Figure 1D). Using a known STAT3 degrader – SD36^28^, we specifically and temporarily removed STAT3, without affecting STAT1 levels (Supplementary Figure 1C). We also measured the dynamics of STAT3 recovery after washout of SD36. We observed that after 3 days without the degrader, STAT3 levels returned close to baseline (Supplementary Figure 1D). With this experimental procedure, we found that STAT3 removal during memory initiation enhanced GBP5 expression in reinduction (Figure 1E), while its removal during recall resulted in a strong reduction of GBP5 level (Figure 1F).

To investigate how STAT3 inhibits IFNγ memory establishment, we examined STAT1 phosphorylation kinetics upon IFNγ stimulation in the function of STAT3 depletion. Removal of STAT3 did not affect the primary STAT1 Tyr701 phosphorylation but impacted the secondary STAT1 Ser727 phosphorylation dynamics (Figure 2A). Additionally, we observed a similar feature in THP-1 macrophages with two different SD36 concentrations: low 1µM and high 10µM. At both SD36 concentrations we observed enhanced STAT1 Ser727 phosphorylation (Supplementary Figure 2A).

**Figure 2.**
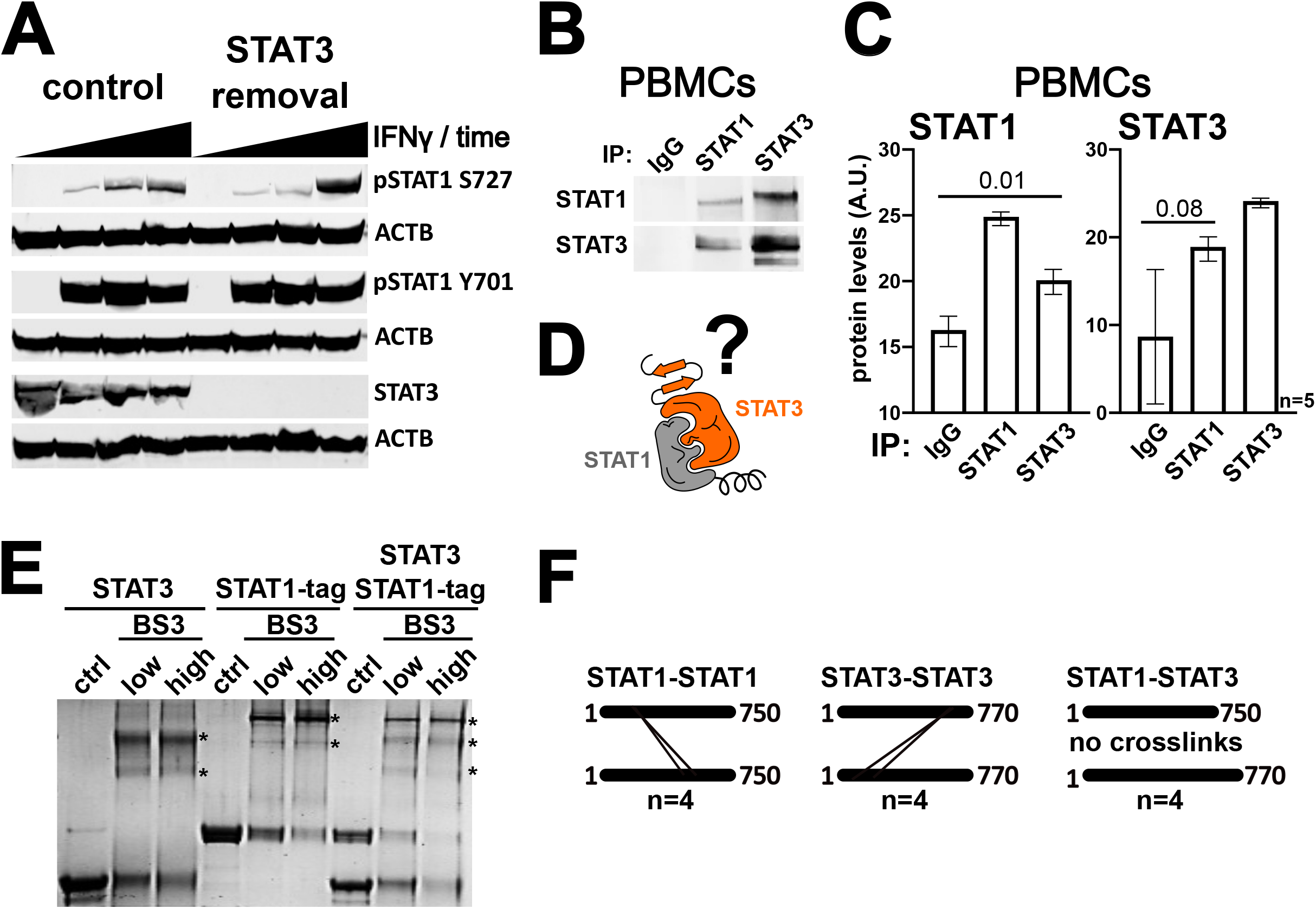
STAT3 indirectly inhibits the initiation of IFNγ memory by reducing STAT1 phosphorylation at serine 727. **(A)** Western blot results showing the effect of STAT3 removal on STAT1 phosphorylation at positions S727 and Y701 as well as STAT1 and STAT3 total protein levels upon treatment with IFNγ in the HeLa cell line at different time points. ACTB – loading control. **(B)** Western blot immunoprecipitation results for STAT1, STAT3 and IgG control in PBMCs. **(C)** As in panel B, but with a quantitative measurement performed via mass spectrometry. **(D)** Graphical representation of STAT1 and STAT3 heterodimerization. **(E)** Coomassie stained SDS-PAGE results for a BS3 mediated crosslinking (low: 0.5µM and high: 1µM BS3 concentration) experiment for STAT1 and STAT3 proteins expressed and purified from *Escherichia coli*. Measurement performed for STAT3 alone (left columns), tagged STAT1 alone (middle columns) and for STAT1-STAT3 mixture (right columns). **(F)** Representation of mass spectrometry results showing crosslinked peptides in STAT1 and STAT3 homodimers, as well as lack of detected interaction for the STAT1-STAT3 mixture.

We hypothesized that STAT3 might directly inhibit STAT1 Ser727 phosphorylation. To test this we performed immunoprecipitation experiments from donor peripheral blood mononuclear cells (PBMCs) followed by western blot and mass spectrometry analysis (MS). This revealed a specific interaction between STAT1 and STAT3 (Figures 2B and 2C). To further test if STAT1-STAT3 is a direct or an indirect interaction (Figure 2D), we produced and purified STAT1 and STAT3 in full length from an *Escherichia coli* heterologous system with nickel and glutathione S-transferase (GST) affinity purification (Supplementary Figure 2B). To asses if the reconstituted proteins are properly folded, we performed calibrated, analytical size exclusion chromatography. The results showed that STAT1 is 139 kDa and STAT3 is 140 kDa, significantly more than their monomeric theoretical masses of 87 kDa and 88 kDa, respectively (Supplementary Figure 2C).

However, as the proteins are of similar size, the analytical size exclusion chromatography could not detect the STAT1-STAT3 heterodimer. To determine if STAT1 and STAT3 can form a direct complex, we performed bissulfosuccinimidyl suberate (BS3)-mediated crosslinking of isolated proteins in separation and mixed with each other in stochiometric proportions. To distinguish between the transcription factors, we used STAT1 fused to the GST-tag and native STAT3. We observed a clear shift in size after crosslink addition resulting in two additional bands for STAT3 and two for STAT1-tag, confirming that the proteins form dimers. However, for the mixed proteins we did not observe any additional bands, suggesting that there is no direct heterodimer formation of STAT1 and STAT3 in their non-phosphorylated forms (Figure 2E). Follow up orthogonal validation via mass spectrometry analysis of non-tagged proteins identified specific homodimer crosslinks for both STAT1 and STAT3, but no heterodimers (Figure 2F). Additionally, with this measurement we were able to detect both intra- and intermolecular crosslinks of STAT3. Interestingly, for the latter, two mutually exclusive crosslinks pointing to coexisting conformations of the STAT3 dimer (Supplementary Figure 2D).

Our study shows that STAT3 has a dual role in regulating IFNγ memory, inhibiting initiation while being necessary for recall. We found that STAT3 removal indirectly impacts STAT1 phosphorylation dynamics but does not form a direct complex with STAT1 in the unphosphorylated state.

### IFNγ memory exhibits sex-bias and stabilization of gene expression upon reinduction

We observed individual variability in IFNγ memory among primary macrophages, with sex-specific differences and stabilization of gene expression during reinduction.

To optimize our measurements in primary macrophages, we analyzed the effects of IFNγ on CD14+ monocytes, within the PBMCs, using flow cytometry. We found that 2 ng/mL of IFNγ induces mild expression of MHC class II (HLA-DRA, a known IFNγ memory gene^11^) and costimulatory factors (CD86 and CD80)^29^, allowing for potential enhancement in the reinduction stage (Supplementary Figures 3A). We confirmed this observation in isolated monocytes (Supplementary Figure 3B).

Using the established conditions, we performed an IFNγ memory experiment followed by an mRNA-seq measurement on primary macrophages differentiated with M-CSF from CD14+ monocytes of male (n=3) and female (n=3) donors (Figure 3A). Comparing reinduced versus induced cells revealed a strong memory effect in male but not female donors, with genes encoding GVINP1 (GTPase, very large interferon inducible pseudogene 1) and B7-H6 (B7 homolog 6) being the most prominent (Figure 3B & Supplementary Table 1). These genes showed stronger reinduction compared to induction in males, but not a classic memory effect due to lack of significant change between naïve and induction conditions (Supplementary Figure 3C).

**Figure 3.**
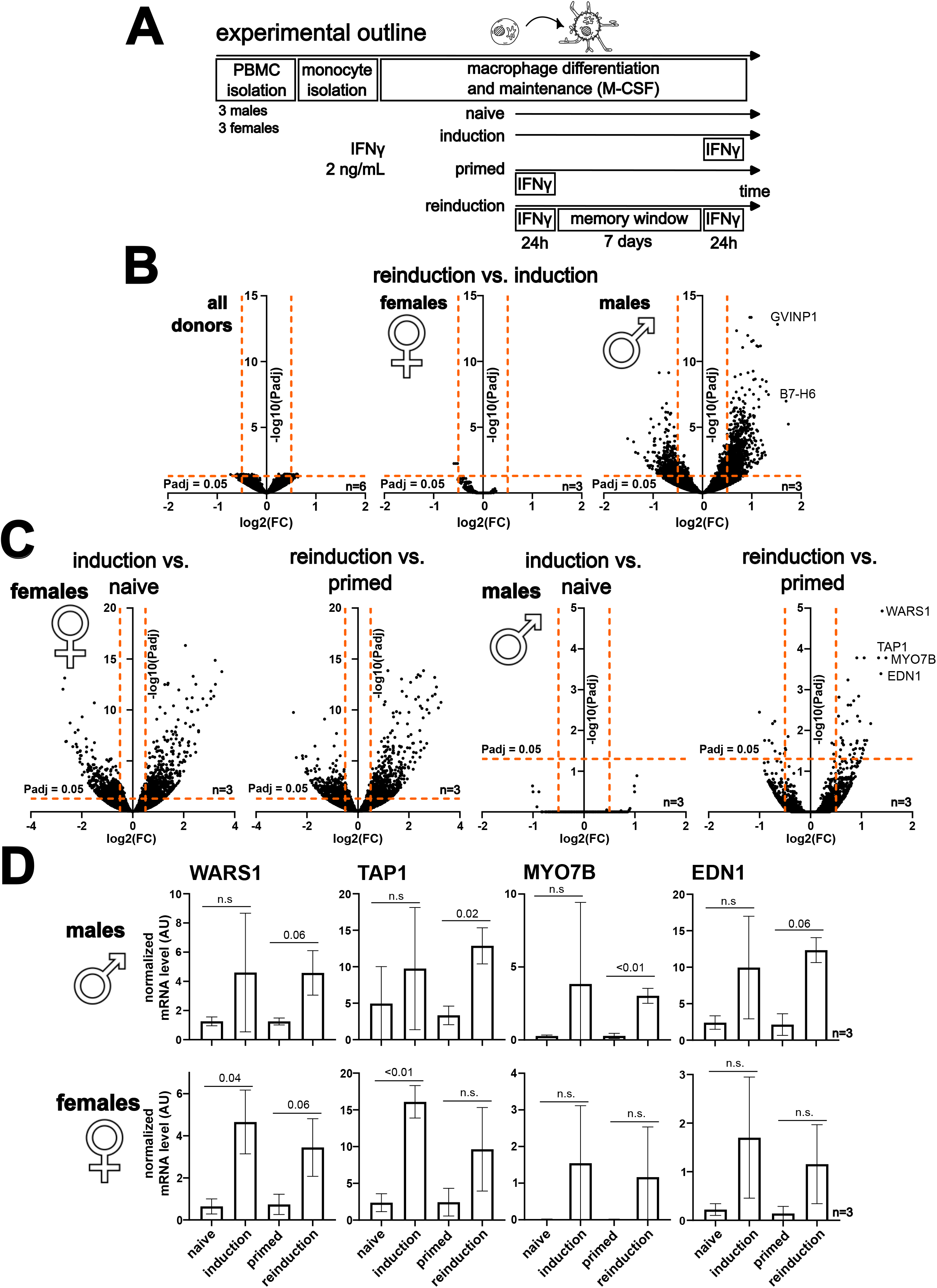
IFNγ memory in human primary macrophages is sex-biased and achieved by reduction of variability in gene expression during reinduction. **(A)** Experimental scheme to measure IFNγ memory in human primary macrophages. **(B)** Results of the experiment from panel A showing differential gene expression in reinduced versus induced primary macrophages for all donors (left), females (middle) and males (right). **(C)** Results of the experiment from panel A showing differential gene expression in induced versus naïve cells and reinduced versus primed cells. Data shown separately for female (left) and male (right) donors. **(D)** Expression level of genes identified in panel C (WARS1, TAP1, MYO7B, EDN1) for different conditions of the IFNγ memory experiment (panel A) in male and female cells.

To better understand the source of this individual variability, we compared induced versus naïve and reinduced versus primed cells separately for females and males. This revealed a strong transcriptome-level difference at the induction stage, primarily due to statistical significance effects, between conditions for male but not female cells (Figure 3C, Supplementary Table 2, 3 & 4). We identified WARS1 (tryptophanyl-tRNA synthetase 1), TAP1 (transporter 1, ATP binding cassette subfamily B member), MYO7B (myosin VIIB) and EDN1 (endothelin 1) as genes with the strongest effect (Figure 3C, Supplementary Table 4). Significant upregulation of these genes relative to induction was observed only during the reinduction step, a feature characteristic for males but absent in females (Figure 3D). We observed the same stabilization of gene expression for the classic memory genes GBP5 and GBP1 (Supplementary Figure 3D). As a next step, we developed a simplified monocyte-to-macrophage differentiation protocol directly from the PBMC population and confirmed its applicability by monitoring cell morphology changes and flow cytometry analysis of monocyte (CD14) and macrophage (CD68) markers (Figure 4A & Supplementary Figure 4.1A). Using this updated protocol, we performed the IFNγ memory experiment followed by an mRNA-seq measurement on isolated PBMCs from new male (n=3) and female (n=3) donors, as in Figure 3, but with a higher IFNγ concentration (50 ng/mL; Figure 4A). As before, males showed a stronger effect than females (Figure 4B & Supplementary Table 5), however the memory outcome was milder than at the lower IFNγ concentration (Figure 3B & Supplementary Table 1). IL32 (interleukin 32) and CCL7 (C-C motif chemokine ligand 7) were identified as top memorized genes, both showing a classic memory effect specifically in male donors (Supplementary Figure 4.1B). Interestingly, CLEC5A (C-type lectin domain containing 5A) showed memory specifically in female cells at the high IFNγ concentration (Figure 4B), but did not represent a classic memory effect due to its high expression in the primed state (Supplementary Figure 4.1C). This makes it likely that the difference between reinduction and induction stages is a result of an elevated expression level in the primed state.

**Figure 4.**
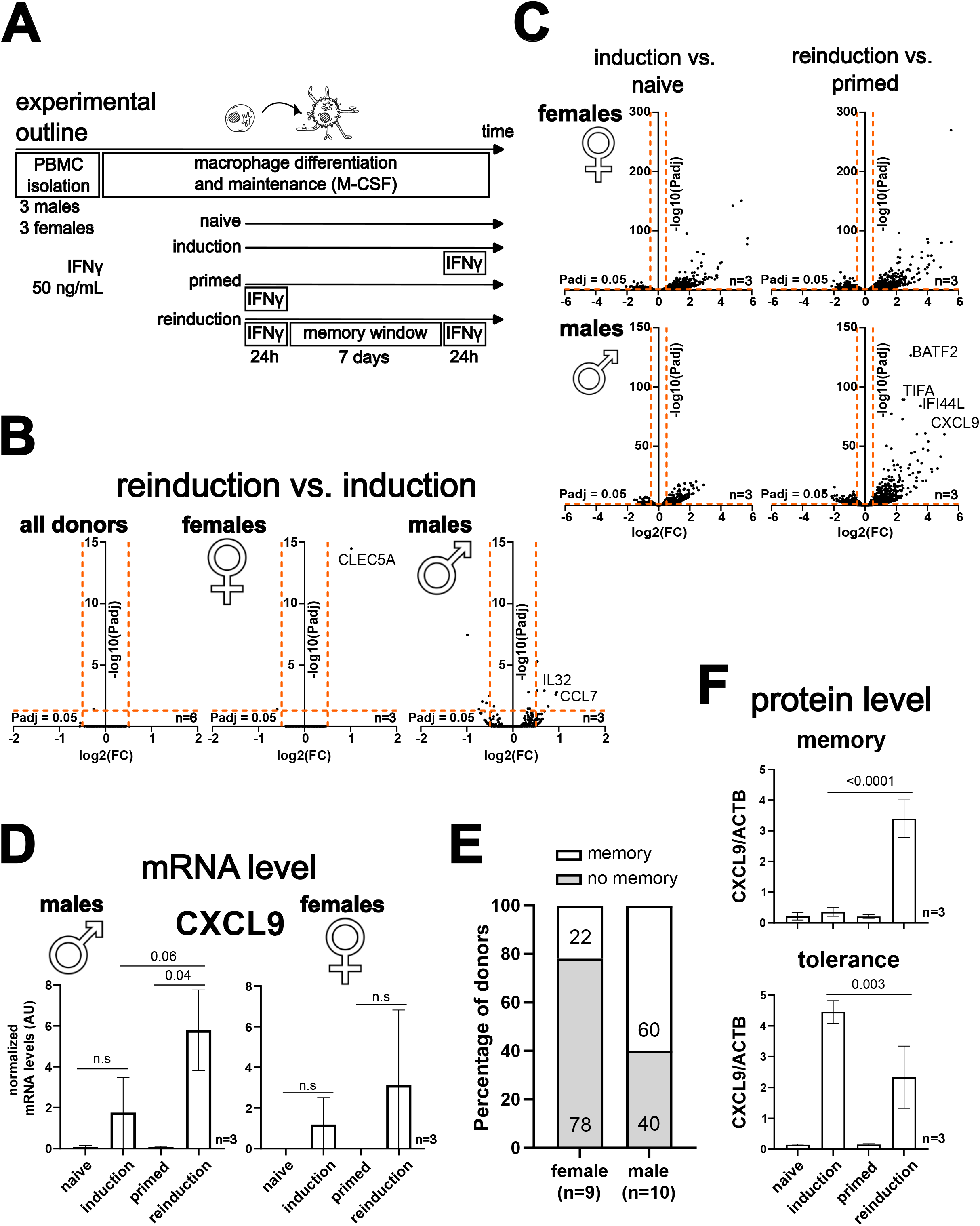
Changes in IFNγ concentration preserve sex-specific effects and reduced gene expression variation during reinduction. **(A)** Simplified experimental scheme to measure IFNγ memory in human primary macrophages, with an additional change of a higher IFNγ concentration (50 ng/mL) than in Figure 3A (2 ng/mL). **(B)** Results of the experiment from panel A showing differential gene expression in reinduced versus induced cells for all donors (left), females (middle) and males (right). **(C)** Results of the experiment from panel A showing differential gene expression in induced versus naïve (left column) and reinduced versus primed (right column) primary macrophages. Data shown separately for female (top row) and male (bottom row) donors. **(D)** Expression level of CXCL9 gene, identified in panel C, in male and female macrophages for different conditions of the IFNγ memory experiment (panel A). **(E)** Percentage of macrophage samples exhibiting or lacking IFNγ memory, derived from female (n = 9) and male (n = 10) donors. **(F)** The representative data from donors whose macrophages showed IFNγ memory (top) or tolerance (bottom).

As previously shown (Figure 3C), comparing induced versus naïve cells and reinduced versus primed cells for females and males separately revealed significant transcriptome-level differences between male and female donors. BATF2 (basic leucine zipper ATF-like transcription factor 2), TIFA (TRAF interacting protein with forkhead associated domain), IFI44L (interferon induced protein 44 like), and CXCL9 (C-X-C motif chemokine ligand 9) were identified as genes with the strongest effect (Figure 4C, Supplementary Table 6, 7, 8 & 9). The reinduction step showed stabilization of gene expression in males but not females, except for IFI44L showing mild reduction of variability in females (Supplementary Figure 4.1D) and CXCL9 showing stabilization and enhanced expression in the reinduction stage specifically in males (Figure 4D).

To further evaluate sex-variability of IFNγ memory, we expanded our analysis to CXCL9 protein expression across a larger cohort of female (n=9) and male (n=10) donors. We used the simplified monocyte-to-macrophage differentiation protocol and a high (50 ng/mL) IFNγ concentration, as under these conditions we observed a memory effect manifested by elevated CXCL9 mRNA levels in male donors (Figure 4D). By performing dot blot measurements for CXCL9 protein, we observed memory in 22% (2 out of 9) female donors and in 60% (6 out of 10) of male donors (Figure 4E, Supplementary Figure 4.2). Surprisingly, aside from memory we also observed tolerance, exclusively in female donors (n=3), marked by lower CXCL9 protein levels in the reinduction in comparison to the induction stage (Figure 4F & Supplementary Figure 4.2).

In conclusion, our results show that IFNγ memory in primary macrophages shows sex-variability with males showing a stronger effect than females; stabilization of gene expression in the reinduction step; and an additional layer of regulation for CXCL9 manifested by tolerance at the protein level.

### IFNγ transcriptional memory is gated by monocyte-to-macrophage differentiation

We established a model system to study IFNγ memory in the context of monocyte and macrophage biology and discovered that cellular differentiation can gate memory recall.

To further investigate the role of IFNγ memory in monocytes and during macrophage differentiation, we developed a model system using the THP-1, male derived, monocyte cell line. This was necessary due to the short lifespan of primary monocytes, with ∼40% dying after one day in culture and more than 60% dead after 8 days (Supplementary Figure 5A). The THP-1 model allows for easy cultivation and rapid differentiation to macrophages^30^. We leveraged these properties to perform the IFNγ memory in THP-1 macrophages (Figure 5A).

**Figure 5.**
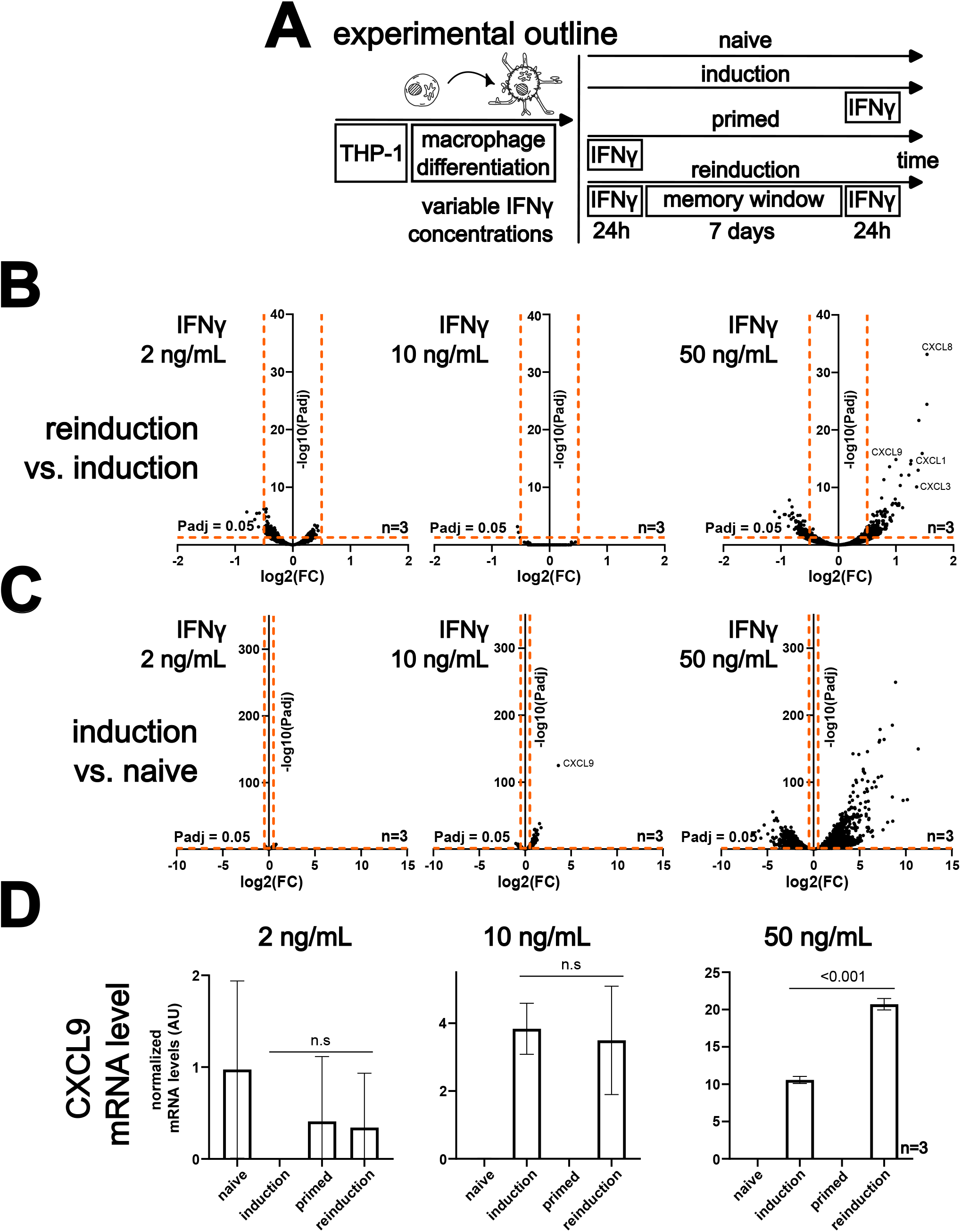
IFNγ memory in THP-1 macrophages is dependent on the cytokine concentration. **(A)** Experimental scheme to measure IFNγ memory in THP-1 macrophages. **(B)** Results of the experiment from panel A showing differential gene expression in reinduced versus induced cells for 2 ng/mL (left), 10 ng/mL (middle) and 50 ng/mL (right) of IFNγ. **(C)** Results of the experiment from panel A showing differential gene expression in induced versus naïve cells at 2 ng/mL (left), 10 ng/mL (middle) and 50 ng/mL (right) of IFNγ. **(D)** Expression levels of CXCL9 gene, identified in panel B, at the indicated IFNγ concentrations and conditions of the IFNγ memory experiment (panel A).

Our experiments with variable IFNγ concentrations in primary cells demonstrated that different genes exhibit memory effects at distinct cytokine amounts. Genes identified at low concentrations did not show memory at high IFNγ concentrations, and vice versa (Figure 3 & 4). Following up on this observation, with our THP-1 model, we tested various IFNγ concentrations (2, 10 and 50 ng/mL) and found that only the highest one (50 ng/mL) induced the memory effect. We identified several genes from the chemokine family: CXCL1, 3, 8 and 9 (Figure 5B & Supplementary Table 10). Notably, CXCL9 was also identified as a memory gene in primary macrophages (Figure 4C, D & Supplementary Table 9). While the middle concentration of 10 ng/mL did not display memory, it did induce a response to the cytokine, with CXCL9 showing clear activation, confirming our previous observation that transcription alone is not sufficient for memory establishment (Figure 5C & Supplementary Table 11)^15^. All identified memory CXCL genes showed significant induction at the middle concentration (10 ng/mL) and clear memory only at the high IFNγ concentration (50 ng/mL) (Figure 5D and Supplementary Figure 5B). To confirm our observations, we also measured and observed CXCL9 memory at the protein level (Supplementary Figure 5C).

Next, using the established THP-1 model system, we investigated whether the memory effect could be detected in cycling monocytes and how monocyte-to-macrophage differentiation affects this phenomenon. We performed experiments with both IFNγ and lipopolysaccharide (LPS), stimulating monocytes, removing the signal, and either letting cells grow for a week before reinduction or differentiating them to macrophages before restimulation (Figure 6A). This measurement revealed memory in monocytes for IFNγ but not LPS, and that memory can withstand differentiation, for both signals (Figure 6B, Supplementary Table 12, 13 & 14).

**Figure 6.**
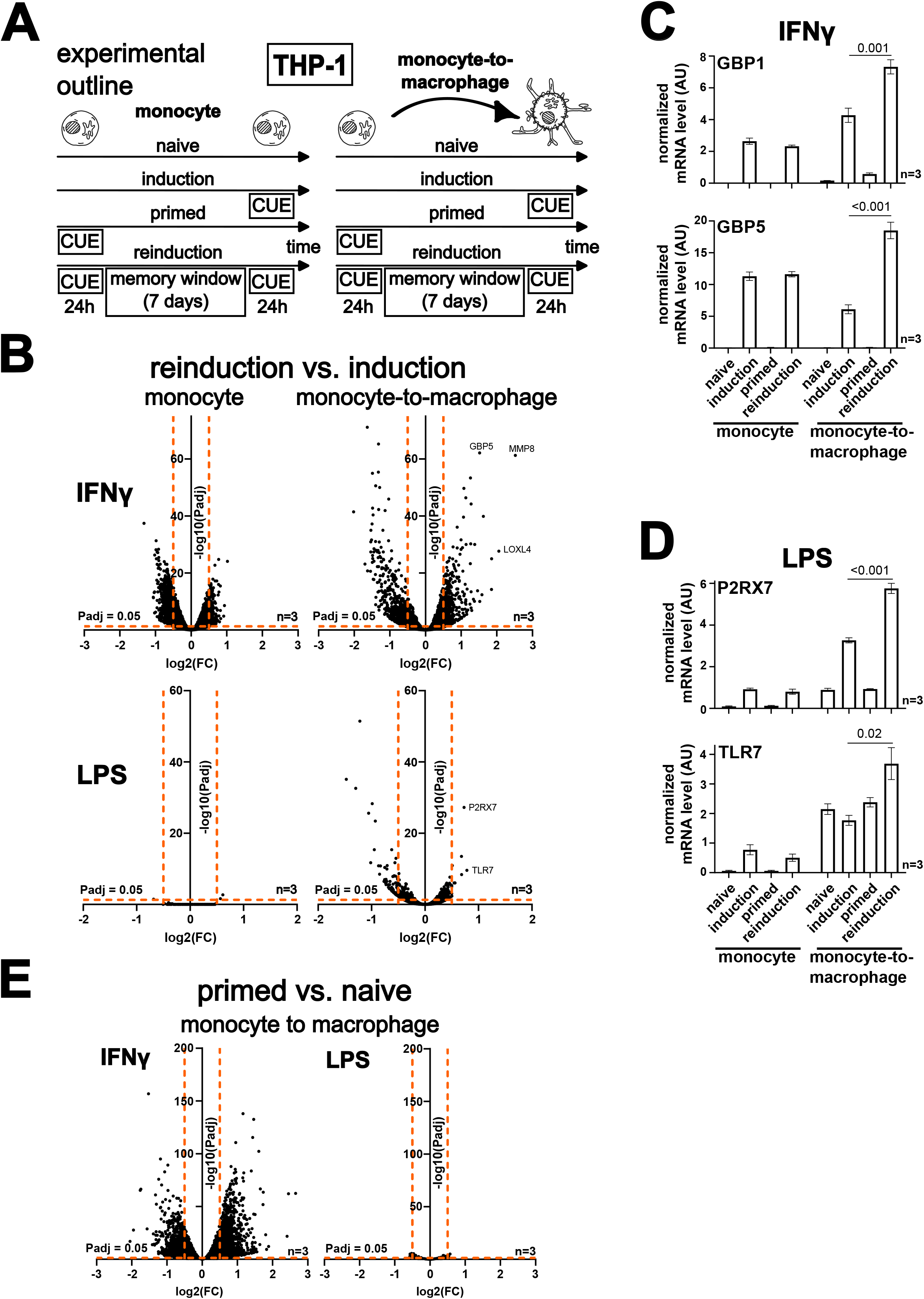
Monocyte-to-macrophage differentiation is required for recall of IFNγ and lipopolysaccharide (LPS) memory. **(A)** Experimental scheme to measure memory with selected cues (IFNγ or LPS) in THP-1 monocytes and during monocyte-to-macrophage differentiation. **(B)** Results of the experiment from panel A showing differential gene expression in reinduced versus induced cells in monocytes (left column) and during monocyte-to-macrophage transition in between the stimulations (right column). Data shown separately for IFNγ (top row) and LPS (bottom row) stimulation. **(C)** Expression levels of GBP1 and GBP5 for different conditions of the IFNγ memory experiment (as in panel A) in monocytes and during monocyte-to-macrophage transition in between the stimulations. **(D)** Expression levels of P2RX7 and TLR7 for different conditions of the LPS memory experiment (as in panel A) in monocytes and during monocyte-to-macrophage transition in the memory window. **(E)** Results of the experiment from panel A showing differential gene expression in primed versus naïve cells during monocyte-to-macrophage transition in the memory window for IFNγ (left panel) and LPS stimulation (right panel).

Interestingly, we observed that classic IFNγ memory genes GBP5 and GBP1 do not show memory in monocytes but exhibit a significant effect during monocyte-to-macrophage transition, indicating that monocytes can remember the exposure but require differentiation for recall of memory for these genes (Figure 6C). We further supported this with an RT-qPCR measurement for GBP5 (Supplementary Figure 6A). Moreover, we expanded this observation to LPS treatment, in which P2RX7 (purinergic receptor P2X 7) and TLR7 (Toll-like receptor 7) genes showed the same differentiation requirement effect (Figure 6B and D).

Additionally, in our data, we identified genes involved in extracellular matrix organization, MMP8 (matrix metallopeptidase 8) and LOXL4 (lysyl oxidase-like 4), which do not display classic IFNγ memory but show higher mRNA levels after induction in the primed state, akin to differentiation (Figure 6B, Supplementary Figure 6B, Supplementary Table 13). To understand this better we compared primed versus naïve cells in the monocyte-to-macrophage memory experiment and revealed that a large number of genes become stably activated by the differentiation signal when primed with IFNγ, but not LPS (Figure 6E & Supplementary Table 15). This suggests that differentiation can gate memory recall through distinct mechanisms.

In conclusion, we developed a model system to study IFNγ memory in monocytes and macrophages. We revealed with it that cellular differentiation plays a critical role in gating memory recall through distinct mechanisms.

## Discussion

This study characterized IFNγ memory in the context of macrophage biology and investigated its underlying mechanisms. IFNγ memory is an important biological process, a part of the broader phenomenon of trained immunity with established functions in health and disease^31^. Our findings indicate that STAT3 plays a dual role in regulating IFNγ memory, inhibiting initiation while being necessary for recall. Additionally, IFNγ memory exhibits sex-variability, achieved by stabilization of gene expression in the reinduction step; with males showing a stronger effect than females. The study also demonstrated that monocyte-to-macrophage differentiation gates memory recall. Overall, these findings provide new insights into the complex regulation of IFNγ memory and its dependence on individual variability and cellular differentiation state.

Our transcriptomic analysis revealed that IFNγ stimulation leads to the overexpression of both STAT1 and STAT3. This finding is consistent with a recent report of IFNγ-induced STAT3 activation in mouse macrophages^32^. Notably, STAT1 and STAT3 often have opposing functions, particularly in cancer, where STAT1 acts as a tumor suppressor by promoting apoptosis and anti-tumor immunity, whereas STAT3 functions as an oncogene that drives cell survival, proliferation, and immune evasion^26^.

We demonstrated that selective removal of STAT3 during the initial induction stage leads to stronger memory, but its removal in the reinduction stage results in impaired memory recall. The effect on memory initiation is a result of STAT3 impacting phosphorylation of STAT1 at position 727. This modification is required for full transcriptional output upon IFNγ stimulation^25^ as well as full-fledged innate immune response against bacterial pathogens^33^. This may be due to the fact that STAT1 and STAT3 are substrates for the same kinase, Cyclin Dependent Kinase 8 (CDK8), a module of the Mediator complex^34^.

Interestingly, we found that STAT1 and STAT3 interact in primary cells without cytokine stimulation, but this interaction is not direct. Reconstitution experiments showed that these proteins do not form heterodimers, consistent with a study in HeLa cells using a co-localization assay^35^. Furthermore, we detected two conformational forms of the unphosphorylated STAT3 homodimer, as previously suggested for this transcription factor^36–38^. As we performed our experiments in a reconstituted system, those forms are directly encoded in the amino acid sequence of STAT3.

Our study of human primary macrophages revealed significant sex-bias in IFNγ memory, with males showing a stronger effect. Sex differences in gene expression programs across various human tissues are well documented^39,40^, but with females exhibiting more reactive and inflammatory immune profiles that contribute to a higher prevalence of autoimmune diseases^41^. At least part of this effect can be explained by a mechanistic study showing that Toll-like receptor 7 (TLR7) escapes X chromosome inactivation in female immune cells, resulting in increased TLR7 protein levels and enhanced responsiveness to TLR7 ligands compared to male cells^42^.

However, the male bias we observed is consistent with clinical reports of sex differences in trained immunity. For example, *Bacillus* Calmette–Guérin (BCG) vaccination in healthy adults showed males tend to exhibit stronger responses, which additionally correlated with baseline testosterone levels^43^. A different, preclinical study on trained immunity to bacterial infection found that female mice had reduced innate immune memory responses compared to males, which was associated with decreased survival from an opportunistic infection caused by *Burkholderia gladioli*. Furthermore, this study demonstrated that progesterone, a female sex hormone, attenuated trained immune responses and that blocking its receptor with an antagonist restored trained immunity in female mice to levels similar to those of males^44^.

We found that male cells displayed stochastic gene expression during initial induction, which became more stable during reinduction, whereas female cells exhibited stable gene expression in both stimulations. This aligns with studies demonstrating sex differences in gene expression, with females showing more robust responses of innate immune genes, particularly for interferon stimulation^45,46^. Additionally, mathematical modeling of the NF-κB regulatory module has shown that stochastic gene regulation can be stabilized by TNF stimulation^47^. The stabilization of gene expression in male cells during reinduction may be regulated by transcriptional bursting, where higher frequency and burst size can stabilize gene expression outputs^48^. While this phenomenon can enable cellular adaptation to changing environments or differentiation into distinct cell types, it can also weaken innate immune responses^49^.

Beyond differences in gene expression, we observed sex differences in the response to IFNγ reinduction at the protein level. While some donors displayed a memory effect, characterized by enhanced CXCL9 protein levels upon secondary stimulation, others exhibited a tolerance phenomenon, marked by a decrease in the secondary response. This tolerogenic response may act as a protective feedback mechanism, preventing excessive pro-inflammatory activation and reducing the risk of tissue damage and autoimmunity^5^.

Our results revealed sex differences in IFNγ memory, with male cells exhibiting stochastic gene expression that stabilizes during reinduction, whereas female cells display stable expression. We also observed differences at the protein level, with some donors showing memory while others exhibiting tolerance for the same gene.

In our work, we observed that different genes exhibit memory in response to varying IFNγ concentrations. This finding is consistent with previous reports of distinct physiological outcomes resulting from different levels of IFNγ stimulation. For example, a low-dose IFNγ has been shown to induce tumor stemness in non-small cell lung cancer, whereas a high-dose IFNγ promotes apoptosis in the same model^50^.

Using the THP-1 system, a well-established tool for studying monocyte and macrophage functions^30^ including in trained immunity^51,52^, we identified a memory effect for the GBP genes and discovered CXCLs (including CXCL9) as novel model genes for studying IFNγ memory in macrophages. Notably, CXCL family genes have been previously shown to exhibit trained immunity phenotypes in response to beta-glucan^53^ and viral models^54^ of training. The regulation of CXCL gene memory involves a long noncoding RNA that recruits the WDR5-MLL1 complex and deposits activating histone marks on gene promoters^53^.

Our study overcame the limitations of primary monocytes – short lifespan, and demonstrated that THP-1 cells can maintain long-term memory of previous IFNγ exposure, which persists even after differentiation into macrophages. This finding is supported by a study showing that training primary monocytes with beta-glucan and allowing them to spontaneously differentiate into macrophages results in a memory effect^55^. Moreover, innate immune memory has been shown to extend beyond mature circulating cells, with the reprogramming of hematopoietic stem and progenitor cells in the bone marrow playing a key role^56^.

Moreover, our work reveals that monocyte-to-macrophage differentiation has a profound impact on memory recall, with certain genes requiring this process for memory retrieval (e.g., GBP5 and GBP1) and others being sufficient for memory recall (e.g., MMP8, LOXL4 and others). This is consistent with a study demonstrating that monocyte-to-macrophage differentiation involves extensive transcriptional and epigenetic changes, including the remodeling of thousands of distal regulatory elements through alterations in histone marks^4^.

In summary, our study reveals that different genes exhibit memory in response to varying IFNγ concentrations and that monocyte-to-macrophage differentiation plays a critical role in gating memory recall. The THP-1 cell line provides a valuable tool for investigating the complex relationships between IFNγ stimulation, gene expression, and immune cell differentiation.

This study provides novel insights into the complex regulation of IFNγ memory, highlighting the dual role of STAT3, sex-specific variability, stabilization of gene expression upon reinduction and the critical impact of monocyte-to-macrophage differentiation on memory recall, ultimately contributing to a deeper understanding of trained immunity, development of personalized therapies and the design of clinical trials.

## Materials and Methods

### Cell culture

Cell lines were maintained at 37°C and 5% CO_2_ in humidified atmosphere. HeLa Kyoto and THP-1 cell lines were cultured in DMEM (Dulbecco’s modified Eagle’s medium) and RPMI (Roswell Park Memorial Institute) 1640 medium, respectively, 10% fetal bovine serum (FBS) and 1% penicillin-streptomycin solution. Cell cultures were regularly screened for mycoplasma contamination using MycoAlert Mycoplasma Detection Kit. Primary immune cells, including PBMCs, primary monocytes, and macrophages were cultured in RPMI 1640 medium containing 20% of FBS and 1% of penicillin-streptomycin solution.

### SD36 PROTAC experiments in HeLa cells

To investigate the role of STAT3 in IFNγ signaling and transcriptional memory, we employed the STAT3-specific PROTAC – SD36^28^. The SD36 stock solution was dissolved in DMSO at concentration of 1 mM and stored at -80°C. Cells were treated with SD36 at a final concentration of 10µM to achieve transient protein depletion, which was validated via Western Blot analysis. Control cells received an equivalent concentration of vehicle (DMSO).

To evaluate the requirement of STAT3 in transcriptional memory, we established an IFNγ priming model combined with stage-specific STAT3 depletion (Figure 1D).

Cells were divided into two parallel cultures: naïve (a mock-treated with medium only) and primed (treated with 50 ng/mL IFNγ for 24 h). Following this treatment, to completely terminate primary signaling, cells were washed with phosphate-buffered saline (PBS) and detached with trypsin solution to remove residual IFNγ and cleave the extracellular domains of membrane receptors^16^. The cells were then maintained in culture for total of 7 days, but on day 5 were detached and seeded in concentration of 3 × 10^4^ cells per well onto 96-well plates to further enable Dot Blot analysis. Following this rest period, both naïve and primed cells were rechallenged with 50 ng/mL of IFNγ for 24 h, and after this time were washed with PBS and processes for Dot Blot analysis.

To selectively assess the role of STAT3 during different phases of transcriptional memory, SD36 treatments were applied at two distinct stages. For the initiation phase, cells were pretreated with SD36 for 24 h prior to and throughout the initial 24 h IFNγ priming. For the recall phase, cells underwent standard priming and the 7-day rest period, followed by SD36 treatment initiated 24 h prior to and maintained throughout the final 24 h IFNγ reinduction.

### Time-course experiments in HeLa cells

To analyze the kinetics of STAT3 depletion, its subsequent impact on the STAT1 expression and its phosphorylation status, a time-course experiment was performed. Cells were seeded onto T25 flask (5 × 10^5^ cell per flask) and the next day were pre-treated with 1 or 10µM SD36 for 24 hours to achieve initial STAT3 depletion. Following the pre-treatment, IFNγ (50 ng/mL) was added to the media to stimulate the cells for 24 h, 6 h, and 3 h prior to harvest, while maintaining continuous exposure to the SD36 throughout the entire duration of the experiment. A vehicle-treated control (DMSO) was maintained under identical conditions. The analysis of target proteins expression (STAT1, pSTAT1(Y701 and S727), STAT3, ACTB – loading control) was performed via Western Blot.

### Western Blot and Dot Blot analyses

Cell lysates for Western Blot analysis were prepared from cell pellets containing up to 8 × 10^6^ cells resuspended in PBS and lysed in Laemmli buffer (125 mM Tris-HCl, pH 6.8, 10% glycerol, 1% sodium dodecyl sulfate (SDS), 0.2% bromophenol blue, and 5% β-mercaptoethanol) at 95°C for 5 minutes. To reduce sample viscosity, lysates were digested with viscolase (500 U/mL). Total nucleic acid concentrations, measured at 260 nm using a NanoDrop Ultra spectrophotometer, served as a proxy for protein content to facilitate sample normalization.

Equal amounts of protein were resolved by SDS-polyacrylamide gel electrophoresis (SDS-PAGE) using 4-12% polyacrylamide gradient gels in Tris-MOPS buffer and transferred to nitrocellulose membranes via wet transfer (100 V, 1 hour, with cooling). Membranes were blocked in 5% non-fat milk in Tris-buffered saline (TBS) containing 0.1% Tween 20 (TBS-T) for 1h at RT. Subsequently, membranes were incubated for 2 h at RT or overnight at 4°C with primary antibodies diluted in 5% milk in TBS-T (for target protein – 1:1,000, for ACTB – 1:2,000 dilution). After washing in TBS-T, the membranes were incubated with fluorophore-conjugated secondary antibodies (1:10,000) for 1h at RT. Fluorescent signals were captured using a Azure 500Q scanner (Azure Biosystems).

Cell lysates for Dot Blot analysis were prepared from approximately 2 – 3 × 10^5^ cells per well in a 96-well plate. Cells were washed with PBS and lysed in a lysis buffer containing 1% Triton X-100, 2 mM MgCl, 0.1% SDS, and 500 U/mL of viscolase in PBS for 15 min. The resulting lysates were transferred onto a nitrocellulose membrane using a Bio-Dot Microfiltration System under vacuum. Subsequently, the membrane was blocked and immunolabeled following the Western Blot procedure described above.

### Immunoprecipitation and sample preparation for mass spectrometry

For immunoprecipitation (IP) and subsequent mass spectrometry analysis, 3 × 10^7^ PBMCs were resuspended in 500 μL of IP buffer (1% n-Dodecyl β-D-maltoside, 150 mM NaCl, 25 mM Tris-HCl, pH 7.5, cOmplete EDTA-Free protease inhibitors, and Viscolase (50 U/mL)). Samples were divided into three groups: non-specific IgG (control), anti-STAT1, and anti-STAT3, each consisting of five replicates. Cell lysis was performed on ice for 30 minutes, and lysates were clarified by centrifugation (6000 × g for 10 minutes at 4°C). Supernatants were transferred to new tubes, and protein concentrations were quantified using a NanoDrop spectrophotometer. Samples were normalized and 500 μg of supernatants were mixed with 5 μg of non-specific IgG, anti-STAT1, or anti-STAT3 antibodies. The mixtures were incubated overnight at 4°C. The following day, Dynabeads Protein A (30 mg/mL) were washed with the IP buffer and added to the lysate antibody mixtures for 45 minutes at RT. The beads were subsequently washed twice with IP buffer and twice with PBS.

To prepare samples for mass spectrometry, immunoprecipitated proteins were eluted in 5 μL of 1M Tris-HCl (pH 8.0) and reduced with 1 μL of 1M dithiothreitol (DTT) for 30 minutes. Alkylation was performed by adding iodoacetamide to a final concentration of 20 mM. Samples were then subjected to overnight digestion with 20 μg of trypsin in 140 μL PBS at 37°C. Finally, supernatants were collected and acidified with trifluoroacetic acid (TFA) to a final concentration of 0.1% prior to mass spectrometry analysis.

### Mass spectrometry measurements

IP samples were separated using a 60-minute gradient on UltiMate™ 3000 RSLC Nano System Flow Meter coupled to Orbitrap Exploris™ 480 Mass Spectrometer. Peptides were concentrated on an Acclaim PepMap 100 C18, 5µm, 5x1mm trap column and separated on C18 PepMap RSLC column (250mm x 0.075mm, 2µm). The mobile phase consisted of 0.1% formic acid in water (A) and 80% acetonitrile with 0.1% formic acid in water (B). Peptides were eluted using a gradient from 2.5% to 40% B.

For cross-linking samples, peptides were separated using a 30-minute gradient on a Vanquish Neo UHPLC System coupled to an Orbitrap Exploris™ 480 Mass Spectrometer. The mobile phase and peptide elution were similar to those used for the IP samples.

Data-dependent (DDA) and data-independent (DIA) acquisitions were collected for IP samples, whereas crosslinking data were DDA-only. All raw files were converted to .mzML files using MSConvert with centroiding.

For the IP experiment, the resulting files were searched against the complete *Homo sapiens* UniProt database (SwissProt + Trembl, all isoforms) concatenated with decoy reverse sequences and contaminants using MSFragger^57^. Search settings included a precursor mass tolerance of +/-10 ppm, automatic fragment mass tolerance optimization, trypsin digestion and peptide length range from 7 to 50 amino acids. MSBooster^58^ was used to re-score results and enhance identification quality.

Quantification was performed using DIA-NN^59^. A spectral library was created *in silico* using a FASTA file identical to that used for the DDA search. The library included spectra for tryptic peptides with maximum one missed cleavage, length range from 7 to 30 amino acids, and methionine oxidation allowed as a variable modification. It was concatenated with experimental library derived from MSFragger search, and library prepared from MSFragger search of pseudo-DDA spectra extracted using DIAUmpire (embedded into FragPipe suite). Merged library were used for quantification.

Cross-linked peptides were identified using MeroX^60^ with subsequent settings. The search database consisted of expected sequences of cross-linked proteins, considering peptide lengths from 5 to 30 amino acids, maximum three missed cleavages, and a cross-linker set as DSS/BS3. Results were exported for visualization using the xiView server^61^.

### STAT1, STAT3 gene synthesis, cloning and construct description

The genes encoding HIS-throm-GST-Press-STAT1 and HIS-throm-GST-Press-STAT3 were codon optimized for *Escherichia coli* expression, synthesized and cloned into the pET24a vector (harboring a kanamycin resistance gene) by GeneScript. The resulting constructs were designed to express fusion proteins with the following components: a histidine (HIS) tag, a thrombin cleavage site, a GST tag, a PreScission Protease cleavage site and STAT1 or STAT3 sequence (Supplementary Table 17).

### Protein expression and purification

BL21 (DE3) *Escherichia coli* were transformed with either the pET24a-HIS-throm-GST-Press-STAT1 or the pET24a-HIS-throm-GST-Press-STAT3 construct. A 2 mL overnight culture was added to 1 L of Luria Broth (LB) medium and grown at 37°C, 230 rpm until an optical density of 0.8 was reached, as measured using an Orion AquaMate 8000 UV-Vis spectrophotometer. The STAT1 construct culture was induced with 0.1 mM isopropyl β-D-1-thiogalactopyranoside (IPTG) and grown overnight at 20°C, 250 rpm, while the STAT3 culture was induced with 1 mM IPTG under the same conditions. Cells were harvested by centrifugation at 6000 × g for 10 minutes at 4°C and stored at -80°C.

For purification, cells were thawed on ice and resuspended in buffer A (50 mM 4-(2-hydroxyethyl)-1-piperazineethanesulfonic acid (HEPES), pH 7.5, 100 mM NaCl, 10% glycerol, 1 mM DTT, 30 mM imidazole, and 0.1% Triton X-100) for 30 minutes. Lysozyme (1 mg/mL) and phenylmethylsulfonyl fluoride (PMSF, 100 μM) were added, and the mixture was incubated on ice for 30 minutes. Lysis was continued using a Vibra Cell Sonicator, and soluble proteins were separated from cell debris by centrifugation at 17,500 × g for 30 minutes at 4°C.

The STAT1-tag and STAT3-tag fusion proteins were purified using a two-step affinity chromatography approach. First, the proteins were bound to a His Trap HP 5 mL column, washed with buffer A, and eluted with buffer B (50 mM HEPES, pH 7.5, 100 mM NaCl, 10% glycerol, 1 mM DTT, 200 mM imidazole, and 0.1% Triton X-100). The eluate was further purified on a GST Trap FF 5 mL column, washed with buffer C (50 mM HEPES, pH 7.5, 100 mM NaCl, 10% glycerol, 1 mM DTT), and eluted with buffer D (50 mM HEPES, pH 7.5, 100 mM NaCl, 10% glycerol, 20 mM glutathione, 1 mM DTT).

For native STAT1 and STAT3 purification, an additional step was included. After binding to the GST Trap column, the proteins were incubated overnight at 8°C with GST-PreScission Protease (0.02 mg/mL) to cleave the tag. The resulting native proteins were then eluted with buffer C from the column.

### Analytical size exclusion chromatography

Size exclusion chromatography was performed on a Superose 6 Increase 5/150 GL column using an ÄKTA pure system. Eluted proteins were detected at 280 nm. To calibrate the column, we used the Gel Filtration Calibration Kit HMW, which contains a mixture of protein standards with known molecular weights: thyroglobulin (669,000 Da), ferritin (440,000 Da), aldolase (158,000 Da), conalbumin (75,000 Da), and ovalbumin (44,000 Da). Blue Dextran 2000 was used to determine the void volume of the column. The mobile phase consisted of 25 mM HEPES, 100 mM NaCl, 10% glycerol, and 1 mM DTT.

The elution points of the calibrant proteins were used to calculate their partition coefficients (K_av_) and then plotted against their logarithmic molecular weights. The elution volumes of STAT1 and STAT3 were used to calculate their K_av_ values, and their molecular weights were determined from the calibration curve. All measurements of elution volumes for calibration proteins and STAT proteins were performed in triplicate.

### BS3 cross linking and sample preparation for mass spectrometry measurement

A 25 mM solution of BS3 cross-linking reagent was prepared by dissolving it in water. Cross-linking reactions were conducted in 40 μL of protein solution at two protein-to-cross-linker molar ratios (1:50 and 1:100) with an incubation time of 40 minutes. The reaction was quenched with 25 mM Tris-HCl, pH 7.5. Cross-linking efficiency was evaluated by observing a shift in band migration during SDS-PAGE.

For mass spectrometry analysis, gel bands representing protein complexes were excised and subjected to a series of washing steps. The gel fragments were washed alternately with buffer E (25% acetonitrile in 37.5 mM ammonium bicarbonate) and buffer F (50% acetonitrile in 25 mM ammonium bicarbonate) at 37°C for 20 minutes, twice. The gel was then reduced with 20 mM DTT in 50 mM ammonium bicarbonate for 30 minutes, followed by alkylation with 60 mM iodoacetamide for 60 minutes in darkness. The gel was washed twice with buffer F for 20 minutes, and the liquid was discarded. The gel was then dehydrated with 100% acetonitrile until it became white.

Trypsin digestion was performed by adding a trypsin solution (10 ng/μL in 50 mM ammonium bicarbonate) to the gel fragments. The samples were incubated overnight at 37°C, and the digestion reaction was quenched with TFA to a final concentration of 0.25%. The samples were sonicated for 15 minutes using an Emmi-D60 sonicator. The supernatant was collected, and acetonitrile was added to the gel fragments, which were then sonicated again for 15 minutes. The combined supernatants were dried in a speedvac (Concentrator Plus).

The resulting protein pellet was dissolved in loading buffer (2.5% acetonitrile, 0.08% TFA) and sonicated. The samples were centrifuged at 13,000 rpm for 10 minutes to remove any remaining debris. The supernatants were processed for mass spectrometry measurements.

### Isolation of Peripheral Blood Mononuclear Cells (PBMCs)

PBMCs were isolated by density gradient centrifugation from buffy coats of healthy donors obtained from the Regional Center for Blood Donation and Blood Treatment in Gdańsk. To isolate PBMCs, 40 mL of diluted blood (1:1 with PBS) was layered over 10 mL of Ficoll-Paque PLUS density gradient medium in a 50 mL tube. Centrifugation was performed at 1,150 × g for 15 minutes at 20°C, with minimal acceleration and no braking.

The resulting PBMC layer was carefully aspirated using a Pasteur pipette and resuspended in PBS to a final volume of 50 mL. The cells were then washed by centrifugation (10 minutes, 415 × g, 20°C) with moderate acceleration and deceleration (acceleration/deceleration = 5). Following washing, the isolated PBMCs were stained with Trypan Blue and counted using TC20 Automated Cell Counter. The cells were then resuspended in cryopreservation medium consisting of 10% DMSO in FBS and stored in liquid nitrogen for subsequent analysis.

### Monocyte isolation

Monocytes were isolated from cryopreserved PBMC fractions using the EasySep Human CD14 Positive Selection Kit II, according to the manufacturer’s protocol. Briefly, thawed PBMCs were resuspended in complete RPMI 1640 medium and centrifuged (5 minutes, 500 × g, 20°C). The cell pellet was then resuspended in EasySep Buffer and the cell number was quantified.

The cells were adjusted to a concentration of 1 × 10^8^ cells/mL, and a maximum volume of 2 mL was transferred to a polystyrene round-bottom tube. The EasySep Human CD14 Positive Selection Cocktail II (200 µL) was added, mixed thoroughly, and incubated for 10 minutes at room temperature (RT). Subsequently, 200 µL of EasySep Dextran RapidSpheres 50100 was added, and the mixture was incubated for an additional 3 minutes at RT.

The suspension was then diluted with 2 mL of EasySep Buffer, mixed, and placed into an EasyEights magnet for 10 minutes at RT. Following incubation, the supernatant was carefully discarded while the tube remained in the magnet. The cell-bead complexes were washed by adding 2.5 mL of EasySep Buffer and the magnet separation was repeated.

Finally, the isolated CD14-positive monocytes were resuspended in the required culture medium for downstream applications.

### Monocyte-to-macrophage differentiation protocols

Isolated monocytes and PBMCs were seeded either into 6-well plates at a density of 2 × 10^6^ cells per well, or into T-25 culture flasks at a maximum density of 2.5 × 10^7^ cells per flask. To induce differentiation into macrophages, the cells were cultured for 7 days in the presence of 100 ng/mL recombinant human macrophage colony-stimulating factor (rhM-CSF). The culture medium with rhM-CSF was replaced on day 4 of the differentiation period. The efficiency of both differentiation protocols was analyzed by flow cytometry by measuring CD14, CD45 and CD68 surface markers (Supplementary Figure 4.1A).

For THP-1 experiments, cells were seeded into T-75 culture flasks at a density of 10^7^ cells per flask in complete RPMI 1640 medium supplemented with 10% FBS. Macrophage differentiation was induced by culturing the cells with 100 nM phorbol 12-myristate-13-acetate (PMA) for 2 days. The medium was then replaced with PMA-free complete RPMI 1640 medium, and the cells were maintained for an additional 2 days prior to any experiments.

### IFNγ memory experiments

To establish memory, cells were split into mock-treated (medium control) and IFNγ-stimulated (50 ng/mL, 24 h) groups. Primary IFNγ stimulation was terminated by washing with and trypsinization. Macrophages intended for dot blot analysis were seeded onto 96-well plates at a maximum density of 3 × 10^5^ cells per well after detachment. Both naive and primed populations were then rested for 7 days in growth medium. Each population was subsequently split and subjected to either a secondary 24 h IFNγ stimulation (50 ng/mL) or mock treatment, yielding four conditions: naive, primed, induced, and reinduced. Finally, cells were processed for dot blot analysis or detached using trypsin, harvested by centrifugation (500 × g, 5 minutes), and the resulting pellets were stored or processed for subsequent analyses.

For the monocyte-to-macrophage memory experiment, THP-1 cells were treated with IFNγ for 24 hours. After priming, the cells were detached and suspended in fresh medium. The cells were then either (1) maintained as monocytes for 7 days or (2) differentiated into macrophages using the PMA protocol described above. Post PMA treatment, the macrophages were washed and maintained in medium for 5 days. Finally, both THP-1 monocytes and macrophages were challenged with IFNγ for 24 hours and harvested for further analysis. In analogous experiments using LPS, a concentration of 100 ng/mL was used.

### Flow cytometry

For flow cytometry analysis, 10^6^ cells were aliquoted per tube and washed twice with 1 mL of PBS via centrifugation (500 × g for 5 minutes) to remove FBS-containing medium. Cells were first incubated with Ghost Dye in the dark for 30 minutes at 4°C for viability assessment according to the manufacturer’s instructions. Following incubation, the cells were washed with PBS containing 1% FBS. To prevent non-specific binding, Fc receptors were blocked by incubating cells with 5 μL of Fc Receptor Blocker in a residual volume of approximately 100 μL for 10 minutes at room temperature. A cocktail of fluorochrome-conjugated antibodies targeting surface markers was then added directly to the cells and incubated for 30 minutes at 4°C in the dark. After staining, the cells were washed with 1 mL of PBS and resuspended in 250 μL of PBS. Data were acquired using a BD FACSAria Fusion Cell Sorter equipped with BD FACSDiva Software and analyzed using FlowJo software.

### RNA extraction, library preparation and sequencing

Primary macrophages or THP-1 cells intended for RNA sequencing were lysed in TRIzol Reagent to ensure RNA stability. Total RNA was extracted from primary macrophages or THP-1 cells. mRNA sequencing libraries were prepared via polyA enrichment. Sequencing was performed on Illumina HiSeq or NovaSeq platforms to generate paired-end 150 bp reads. RNA extraction, library preparation and sequencing were performed by Azenta Life Science or Novogene companies. Subsequently, raw reads in FASTQ format were assessed for quality control using FastQC (v0.12.0). Adapter contamination and low-quality bases were removed with FASTX-Trimmer (v0.0.14) by trimming the first ∼15 bases from both read ends. The trimmed reads were then aligned to the human reference genome (hg38) using bowtie2 (v2.5.4) with local mapping option^62^. PCR duplicates were removed using samtools^63^. Gene expression counts were quantified using htseq-count (v2.1.2)^64^ with the hg38.refGene.gtf annotation file, using union mode, feature type: exon, and ID attribute: gene_id. Differential gene expression analysis was performed using the DESeq2 package (v1.40.2)^65^. DESeq2 was run with a local fit type and beta priors enabled. The results were extracted with an alpha value of 0.05 for the MA-plot. Significance thresholds were set at an adjusted p-value < 0.05 and an absolute log2 fold change > 0.5.

### Reverse Transcription Quantitative Polymerase Chain Reaction (RT-qPCR)

Total RNA was isolated from cells using the InviTrap Spin Universal RNA Mini Kit, according to the manufacturer’s instructions. Reverse transcription was performed on 1 μg of total RNA (quantified by Nanodrop Ultra spectrophotometer) using the Transcriptor First Strand Synthesis Kit and random hexamers.

The expression levels of *GBP5* and the reference gene actin beta (*ACTB*) were analyzed via quantitative PCR (qPCR) using previously described primers^14^ (Supplementary Table 16). qPCR reactions were performed in triplicate on 96-well plates using the PowerUp SYBR Green Master Mix and the CFX96 Real-Time PCR Detection System. The thermal cycling conditions consisted of an initial denaturation at 95°C for 10 minutes, followed by 40 cycles of 95°C for 15 seconds and 60°C for 1 minute. Melting curve analysis was performed from 65 to 95°C with 0.5°C increments to confirm amplicon specificity. To account for run-to-run variation, quantification cycles were corrected. The expression levels of GBP5 were normalized to the reference gene *ACTB* using the ΔΔCt method.

### Statistical analysis

Statistical analyses and data visualization were performed using GraphPad Prism software and Inkscape tool. Continuous variables were summarized as mean ±SD, and differences between groups were assessed via Welch’s t-test. A *P*-value of < 0.05 was considered statistically significant.

## Resources Table

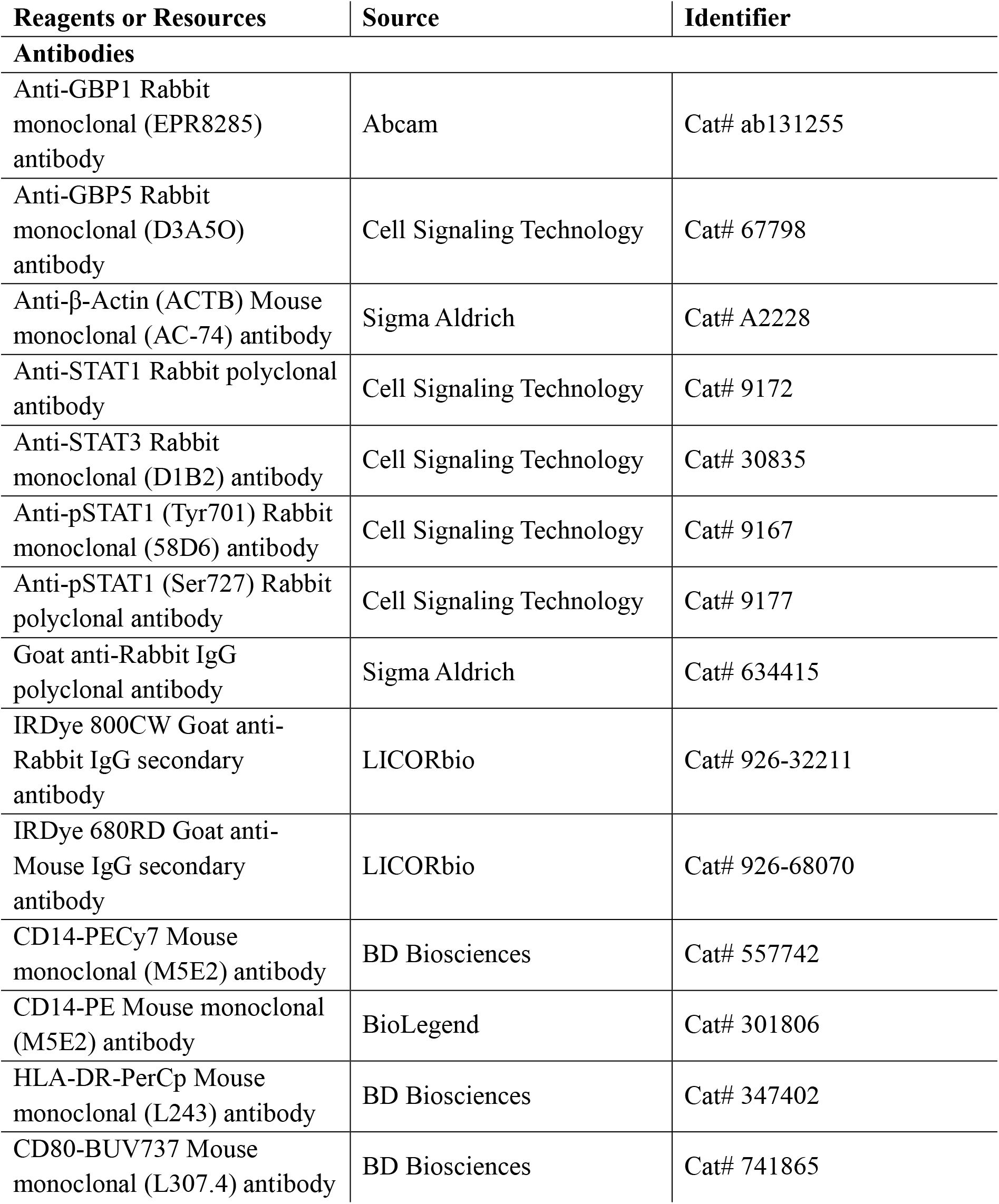
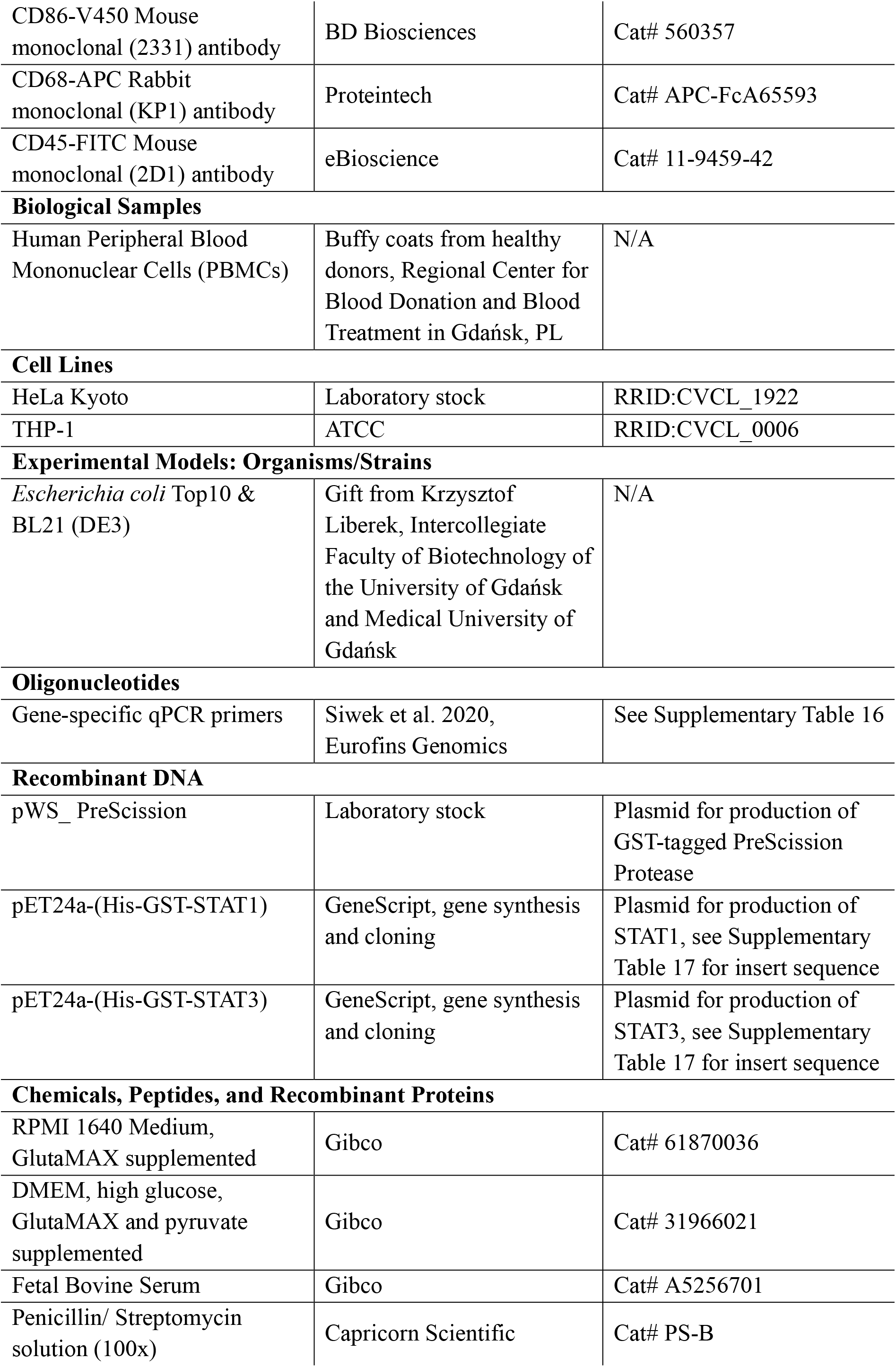
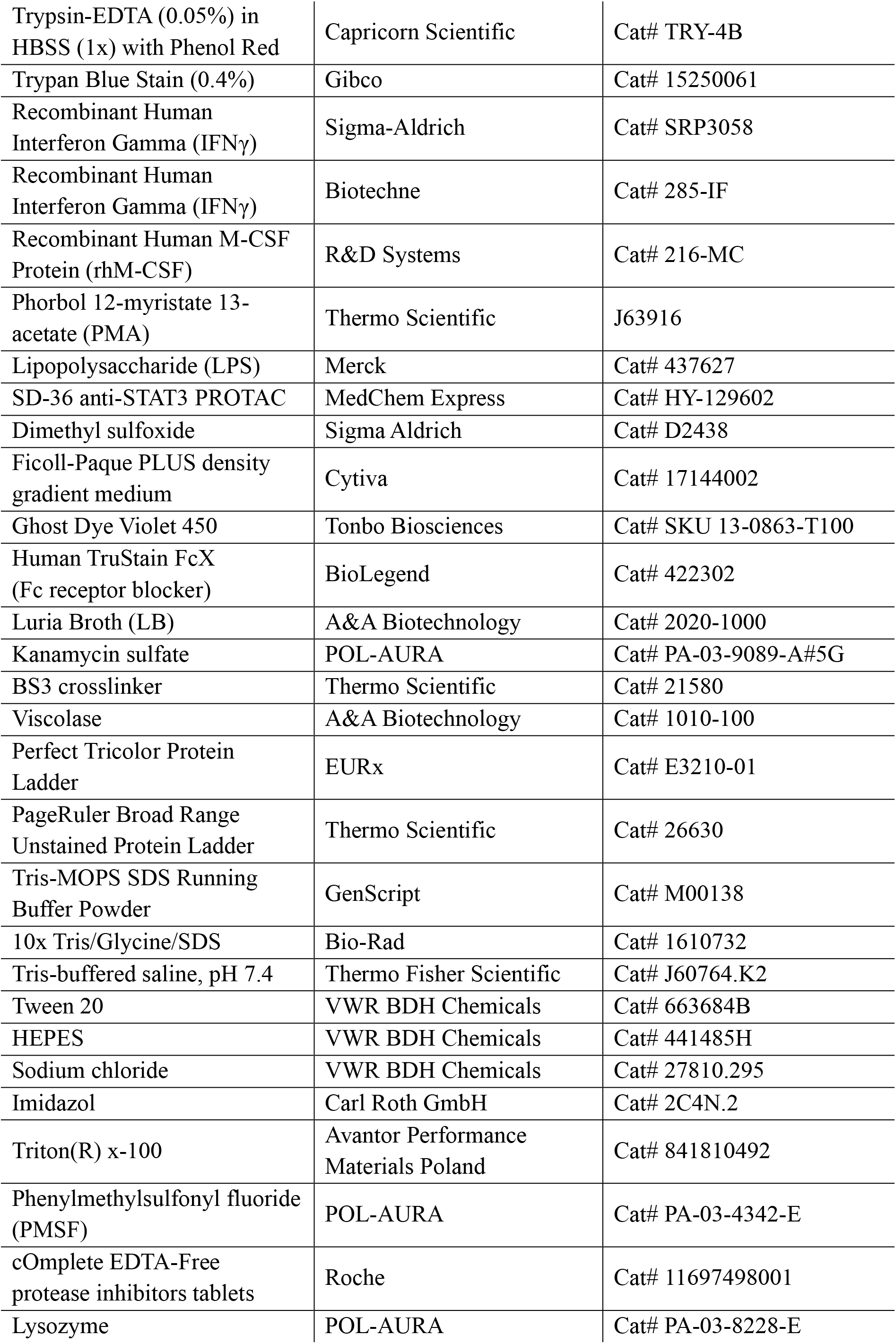
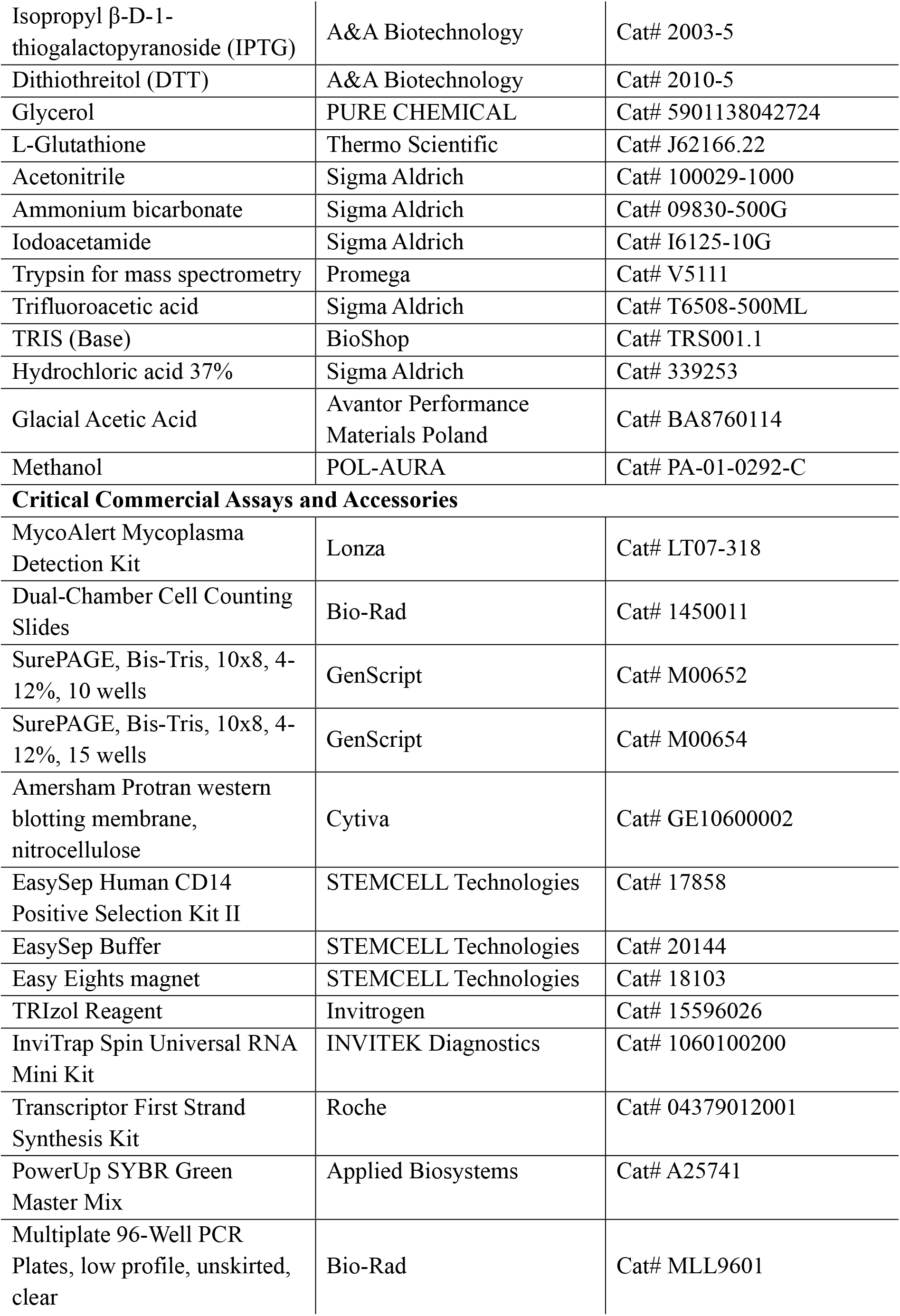
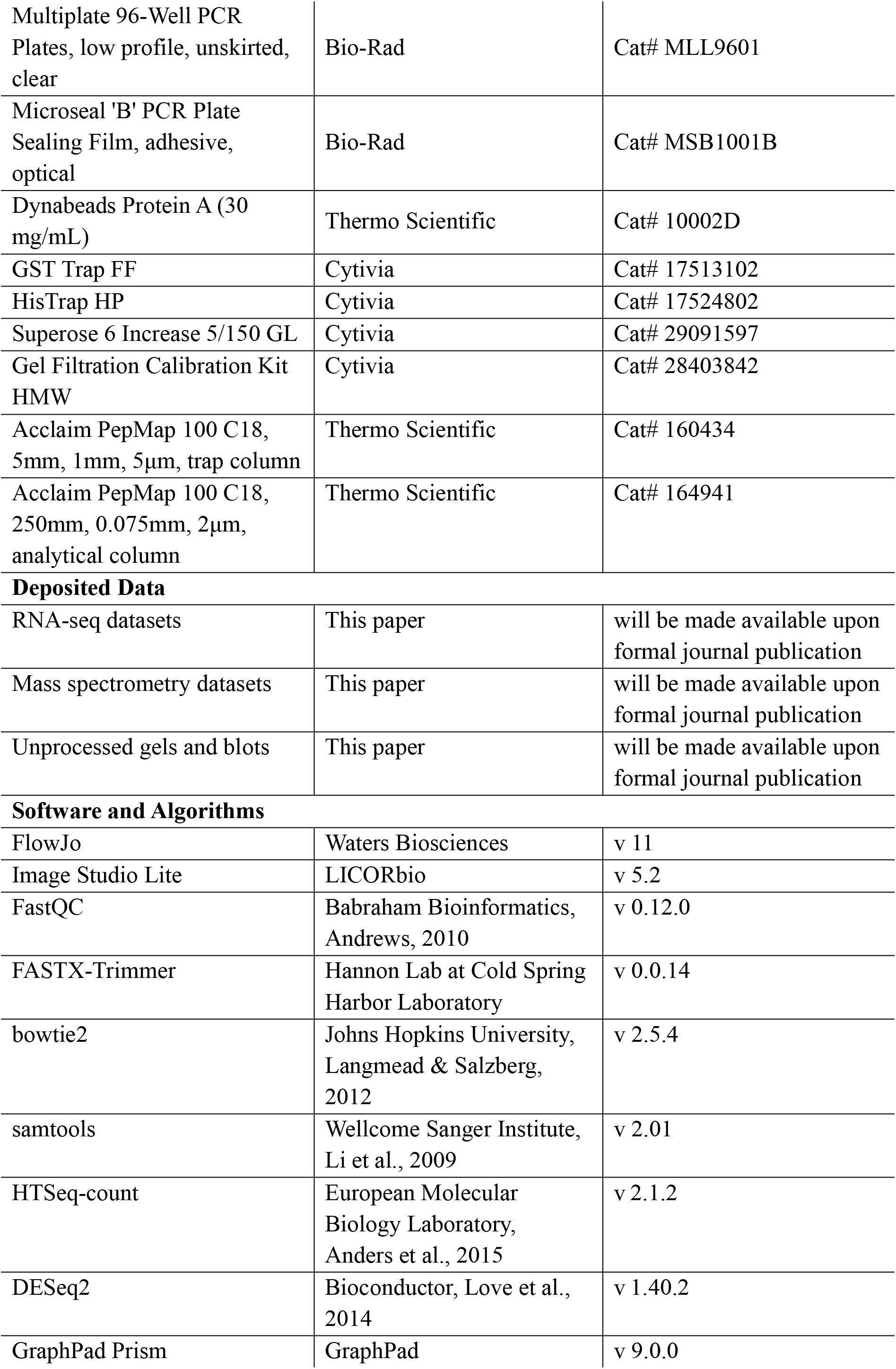
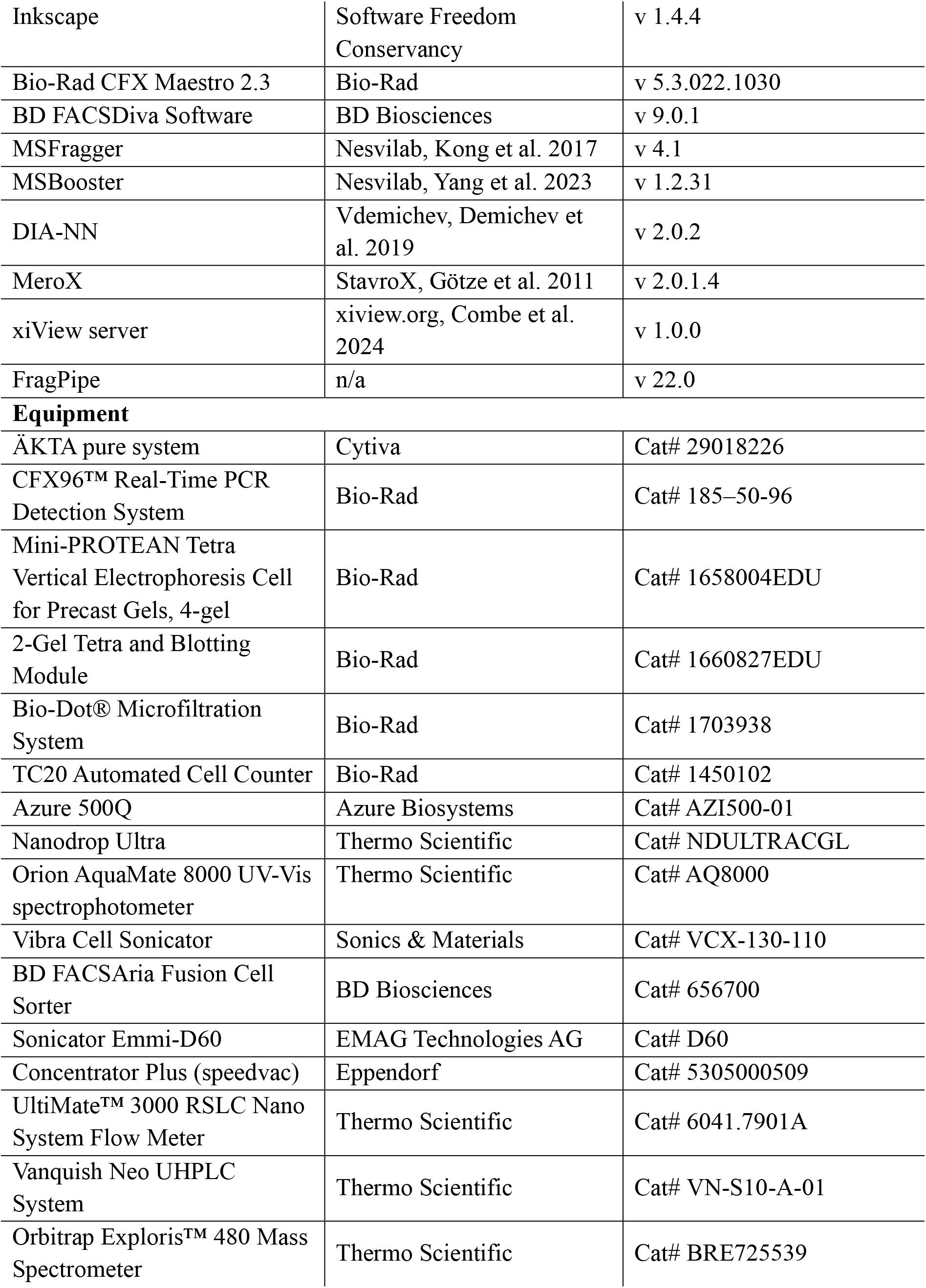

| Reagents or Resources | Source | Identifier |
| --- | --- | --- |
| **Antibodies** |  |  |
| Anti-GBP1 Rabbit monoclonal (EPR8285) antibody | Abcam | Cat# ab131255 |
| Anti-GBP5 Rabbit monoclonal (D3A5O) antibody | Cell Signaling Technology | Cat# 67798 |
| Anti-β-Actin (ACTB) Mouse monoclonal (AC-74) antibody | Sigma Aldrich | Cat# A2228 |
| Anti-STAT1 Rabbit polyclonal antibody | Cell Signaling Technology | Cat# 9172 |
| Anti-STAT3 Rabbit monoclonal (D1B2) antibody | Cell Signaling Technology | Cat# 30835 |
| Anti-pSTAT1 (Tyr701) Rabbit monoclonal (58D6) antibody | Cell Signaling Technology | Cat# 9167 |
| Anti-pSTAT1 (Ser727) Rabbit polyclonal antibody | Cell Signaling Technology | Cat# 9177 |
| Goat anti-Rabbit IgG polyclonal antibody | Sigma Aldrich | Cat# 634415 |
| IRDye 800CW Goat anti-Rabbit IgG secondary antibody | LICORbio | Cat# 926-32211 |
| IRDye 680RD Goat anti-Mouse IgG secondary antibody | LICORbio | Cat# 926-68070 |
| CD14-PECy7 Mouse monoclonal (M5E2) antibody | BD Biosciences | Cat# 557742 |
| CD14-PE Mouse monoclonal (M5E2) antibody | BioLegend | Cat# 301806 |
| HLA-DR-PerCp Mouse monoclonal (L243) antibody | BD Biosciences | Cat# 347402 |
| CD80-BUV737 Mouse monoclonal (L307.4) antibody | BD Biosciences | Cat# 741865 |
| CD86-V450 Mouse monoclonal (2331) antibody | BD Biosciences | Cat# 560357 |
| CD68-APC Rabbit monoclonal (KP1) antibody | Proteintech | Cat# APC-FcA65593 |
| CD45-FITC Mouse monoclonal (2D1) antibody | eBioscience | Cat# 11-9459-42 |
| **Biological Samples** |  |  |
| Human Peripheral Blood Mononuclear Cells (PBMCs) | Buffly coats from healthy donors, Regional Center for Blood Donation and Blood Treatment in Gdańsk, PL | N/A |
| **Cell Lines** |  |  |
| HeLa Kyoto | Laboratory stock | RRID:CVCL_1922 |
| THP-1 | ATCC | RRID:CVCL_0006 |
| **Experimental Models: Organisms/Strains** |  |  |
| *Escherichia coli* Top10 & BL21 (DE3) | Gift from Krzysztof Liberek, Intercollegiate Faculty of Biotechnology of the University of Gdańsk and Medical University of Gdańsk | N/A |
| **Oligonucleotides** |  |  |
| Gene-specific qPCR primers | Siwek et al. 2020, Eurofins Genomics | See Supplementary Table 16 |
| **Recombinant DNA** |  |  |
| pWS_ PreScission | Laboratory stock | Plasmid for production of GST-tagged PreScission Protease |
| pET24a-(His-GST-STAT1) | GeneScript, gene synthesis and cloning | Plasmid for production of STAT1, see Supplementary Table 17 for insert sequence |
| pET24a-(His-GST-STAT3) | GeneScript, gene synthesis and cloning | Plasmid for production of STAT3, see Supplementary Table 17 for insert sequence |
| **Chemicals, Peptides, and Recombinant Proteins** |  |  |
| RPMI 1640 Medium, GlutaMAX supplemented | Gibco | Cat# 61870036 |
| DMEM, high glucose, GlutaMAX and pyruvate supplemented | Gibco | Cat# 31966021 |
| Fetal Bovine Serum | Gibco | Cat# A5256701 |
| Penicillin/ Streptomycin solution (100x) | Capricorn Scientific | Cat# PS-B |
| Trypsin-EDTA (0.05%) in HBSS (1x) with Phenol Red | Capricorn Scientific | Cat# TRY-4B |
| Trypan Blue Stain (0.4%) | Gibco | Cat# 15250061 |
| Recombinant Human Interferon Gamma (IFNγ) | Sigma-Aldrich | Cat# SRP3058 |
| Recombinant Human Interferon Gamma (IFNγ) | Biotechne | Cat# 285-IF |
| Recombinant Human M-CSF Protein (rhM-CSF) | R&D Systems | Cat# 216-MC |
| Phorbol 12-myristate 13-acetate (PMA) | Thermo Scientific | J63916 |
| Lipopolysaccharide (LPS) | Merck | Cat# 437627 |
| SD-36 anti-STAT3 PROTAC | MedChem Express | Cat# HY-129602 |
| Dimethyl sulfoxide | Sigma Aldrich | Cat# D2438 |
| Ficoll-Paque PLUS density gradient medium | Cytiva | Cat# 17144002 |
| Ghost Dye Violet 450 | Tonbo Biosciences | Cat# SKU 13-0863-T100 |
| Human TruStain FcX (Fc receptor blocker) | BioLegend | Cat# 422302 |
| Luria Broth (LB) | A&A Biotechnology | Cat# 2020-1000 |
| Kanamycin sulfate | POL-AURA | Cat# PA-03-9089-A#5G |
| BS3 crosslinker | Thermo Scientific | Cat# 21580 |
| Viscolase | A&A Biotechnology | Cat# 1010-100 |
| Perfect Tricolor Protein Ladder | EURx | Cat# E3210-01 |
| PageRuler Broad Range Unstained Protein Ladder | Thermo Scientific | Cat# 26630 |
| Tris-MOPS SDS Running Buffer Powder | GenScript | Cat# M00138 |
| 10x Tris/Glycine/SDS | Bio-Rad | Cat# 1610732 |
| Tris-buffered saline, pH 7.4 | Thermo Fisher Scientific | Cat# J60764.K2 |
| Tween 20 | VWR BDH Chemicals | Cat# 663684B |
| HEPES | VWR BDH Chemicals | Cat# 441485H |
| Sodium chloride | VWR BDH Chemicals | Cat# 27810.295 |
| Imidazol | Carl Roth GmbH | Cat# 2C4N.2 |
| Triton(R) x-100 | Avantor Performance Materials Poland | Cat# 841810492 |
| Phenylmethylsulfonyl fluoride (PMSF) | POL-AURA | Cat# PA-03-4342-E |
| cOmplete EDTA-Free protease inhibitors tablets | Roche | Cat# 11697498001 |
| Lysozyme | POL-AURA | Cat# PA-03-8228-E |
| Isopropyl β-D-1-thiogalactopyranoside (IPTG) | A&A Biotechnology | Cat# 2003-5 |
| Dithiothreitol (DTT) | A&A Biotechnology | Cat# 2010-5 |
| Glycerol | PURE CHEMICAL | Cat# 5901138042724 |
| L-Glutathione | Thermo Scientific | Cat# J62166.22 |
| Acetonitrile | Sigma Aldrich | Cat# 100029-1000 |
| Ammonium bicarbonate | Sigma Aldrich | Cat# 09830-500G |
| Iodoacetamide | Sigma Aldrich | Cat# I6125-10G |
| Trypsin for mass spectrometry | Promega | Cat# V5111 |
| Trifluoroacetic acid | Sigma Aldrich | Cat# T6508-500ML |
| TRIS (Base) | BioShop | Cat# TRS001.1 |
| Hydrochloric acid 37% | Sigma Aldrich | Cat# 339253 |
| Glacial Acetic Acid | Avantor Performance Materials Poland | Cat# BA8760114 |
| Methanol | POL-AURA | Cat# PA-01-0292-C |
| **Critical Commercial Assays and Accessories** |  |  |
| MycoAlert Mycoplasma Detection Kit | Lonza | Cat# LT07-318 |
| Dual-Chamber Cell Counting Slides | Bio-Rad | Cat# 1450011 |
| SurePAGE, Bis-Tris, 10x8, 4-12%, 10 wells | GenScript | Cat# M00652 |
| SurePAGE, Bis-Tris, 10x8, 4-12%, 15 wells | GenScript | Cat# M00654 |
| Amersham Protran western blotting membrane, nitrocellulose | Cytiva | Cat# GE10600002 |
| EasySep Human CD14 Positive Selection Kit II | STEMCELL Technologies | Cat# 17858 |
| EasySep Buffer | STEMCELL Technologies | Cat# 20144 |
| Easy Eights magnet | STEMCELL Technologies | Cat# 18103 |
| TRIzol Reagent | Invitrogen | Cat# 15596026 |
| InviTrap Spin Universal RNA Mini Kit | INVITEK Diagnostics | Cat# 1060100200 |
| Transcriptor First Strand Synthesis Kit | Roche | Cat# 04379012001 |
| PowerUp SYBR Green Master Mix | Applied Biosystems | Cat# A25741 |
| Multiplate 96-Well PCR Plates, low profile, unskirted, clear | Bio-Rad | Cat# MLL9601 |
| Multiplate 96-Well PCR Plates, low profile, unskirted, clear | Bio-Rad | Cat# MLL9601 |
| Microseal 'B' PCR Plate Sealing Film, adhesive, optical | Bio-Rad | Cat# MSB1001B |
| Dynabeads Protein A (30 mg/mL) | Thermo Scientific | Cat# 10002D |
| GST Trap FF | Cytivia | Cat# 17513102 |
| HisTrap HP | Cytivia | Cat# 17524802 |
| Superose 6 Increase 5/150 GL | Cytivia | Cat# 29091597 |
| Gel Filtration Calibration Kit HMW | Cytivia | Cat# 28403842 |
| Acclaim PepMap 100 C18, 5mm, 1mm, 5µm, trap column | Thermo Scientific | Cat# 160434 |
| Acclaim PepMap 100 C18, 250mm, 0.075mm, 2µm, analytical column | Thermo Scientific | Cat# 164941 |
| **Deposited Data** |  |  |
| RNA-seq datasets | This paper | will be made available upon formal journal publication |
| Mass spectrometry datasets | This paper | will be made available upon formal journal publication |
| Unprocessed gels and blots | This paper | will be made available upon formal journal publication |
| **Software and Algorithms** |  |  |
| FlowJo | Waters Biosciences | v 11 |
| Image Studio Lite | LICORbio | v 5.2 |
| FastQC | Babraham Bioinformatics, Andrews, 2010 | v 0.12.0 |
| FASTX-Trimmer | Hannon Lab at Cold Spring Harbor Laboratory | v 0.0.14 |
| bowtie2 | Johns Hopkins University, Langmead & Salzberg, 2012 | v 2.5.4 |
| samtools | Wellcome Sanger Institute, Li et al., 2009 | v 2.01 |
| HTSeq-count | European Molecular Biology Laboratory, Anders et al., 2015 | v 2.1.2 |
| DESeq2 | Bioconductor, Love et al., 2014 | v 1.40.2 |
| GraphPad Prism | GraphPad | v 9.0.0 |
| Inkscape | Software Freedom Conservancy | v 1.4.4 |
| Bio-Rad CFX Maestro 2.3 | Bio-Rad | v 5.3.022.1030 |
| BD FACSDiva Software | BD Biosciences | v 9.0.1 |
| MSFragger | Nesvilab, Kong et al. 2017 | v 4.1 |
| MSBooster | Nesvilab, Yang et al. 2023 | v 1.2.31 |
| DIA-NN | Vdemichev, Demichev et al. 2019 | v 2.0.2 |
| MeroX | StavroX, Götze et al. 2011 | v 2.0.1.4 |
| xiView server | xiview.org, Combe et al. 2024 | v 1.0.0 |
| FragPipe | n/a | v 22.0 |
| Equipment |  |  |
| ÄKTA pure system | Cytiva | Cat# 29018226 |
| CFX96™ Real-Time PCR Detection System | Bio-Rad | Cat# 185–50-96 |
| Mini-PROTEAN Tetra Vertical Electrophoresis Cell for Precast Gels, 4-gel | Bio-Rad | Cat# 1658004EDU |
| 2-Gel Tetra and Blotting Module | Bio-Rad | Cat# 1660827EDU |
| Bio-Dot® Microfiltration System | Bio-Rad | Cat# 1703938 |
| TC20 Automated Cell Counter | Bio-Rad | Cat# 1450102 |
| Azure 500Q | Azure Biosystems | Cat# AZI500-01 |
| Nanodrop Ultra | Thermo Scientific | Cat# NDULTRACGL |
| Orion AquaMate 8000 UV-Vis spectrophotometer | Thermo Scientific | Cat# AQ8000 |
| Vibra Cell Sonicator | Sonics & Materials | Cat# VCX-130-110 |
| BD FACSAria Fusion Cell Sorter | BD Biosciences | Cat# 656700 |
| Sonicator Emmi-D60 | EMAG Technologies AG | Cat# D60 |
| Concentrator Plus (speedvac) | Eppendorf | Cat# 5305000509 |
| UltiMate™ 3000 RSLC Nano System Flow Meter | Thermo Scientific | Cat# 6041.7901A |
| Vanquish Neo UHPLC System | Thermo Scientific | Cat# VN-S10-A-01 |
| Orbitrap Exploris™ 480 Mass Spectrometer | Thermo Scientific | Cat# BRE725539 |

## Data Availability

The RNA-seq transcriptomics and mass spectrometry proteomics datasets as well as unaltered images of all western blots and dot blots will be made available upon formal journal publication.

## Acknowledgments

W.Si. would like to thank Robert E. Kingston for creating a stimulating, supportive, and intellectually enriching environment in his laboratory during the early stages of this work. We acknowledge Ines Papak and Anna Biernacka for their help with flow cytometry measurements, as well as Iwona Skorowska and Martyna Siewiera for their support with cell culture. We thank Dorota Krzyżanowska for granting us access to an RT-qPCR instrument and Krzysztof Liberek for generously providing bacterial strains used in this study. Additionally, we extend our gratitude to Honorata Czapińska and Anton Slyvka for their helpful comments and feedback, which improved the quality of this work

## Author Contributions

Conceptualization: W.Si.; Methodology: J.T., W.Sn., M.M., S.Ko., A.P., E.C., W.Si.; Software: S.Kh., A.P., W. Si.; Validation: J.T., W.Sn., M.M., A.P., E.C., W.Si.; Formal analysis: J.T., W.Sn., M.M., S.Kh., A.P., E.C., W.Si.; Investigation: J.T., W.Sn., M.M., A.P., E.C., W.Si.; Resources: S.Ko., W.Si.; Data curation: J.T., W.Sn., M.M., S.Kh., A.P., E.C., W.Si.; Writing – Original Draft: J.T., W.Si.; Writing – Review & Editing: J.T., W.Sn., M.M., S.Kh., S.Ko., A.P., E.C., W.Si.; Visualization: J.T., W.Sn., M.M., S.Kh., A.P., W.Si.; Supervision: W.Si.; Project administration: W.Si; Funding acquisition: J.T., S.Ko., W.Si.

## Supplemental Information

Supplementary Document 1. Supplementary Figures 1-6.

Supplementary Table 1. List of genes from the IFNγ memory experiment, male primary macrophages, cytokine concentration 2 ng/mL. Related to Figure 3B.

Supplementary Table 2. List of genes from the IFNγ induction experiment, female primary macrophages, cytokine concentration 2 ng/mL. Related to Figure 3C.

Supplementary Table 3. List of genes from the IFNγ reinduction experiment, female primary macrophages, cytokine concentration 2 ng/mL. Related to Figure 3C.

Supplementary Table 4. List of genes from the IFNγ reinduction experiment, male primary macrophages, cytokine concentration 2 ng/mL. Related to Figure 3C.

Supplementary Table 5. List of genes from the IFNγ memory experiment, male primary macrophages, cytokine concentration 50 ng/mL. Related to Figure 4B.

Supplementary Table 6. List of genes from the IFNγ induction experiment, female primary macrophages, cytokine concentration 50 ng/mL. Related to Figure 4C.

Supplementary Table 7. List of genes from the IFNγ reinduction experiment, female primary macrophages, cytokine concentration 50 ng/mL. Related to Figure 4C.

Supplementary Table 8. List of genes from the IFNγ induction experiment, male primary macrophages, cytokine concentration 50 ng/mL. Related to Figure 4C.

Supplementary Table 9. List of genes from the IFNγ reinduction experiment, male primary macrophages, cytokine concentration 50 ng/mL. Related to Figure 4C.

Supplementary Table 10. List of genes from the IFNγ memory experiment, THP-1 macrophages, cytokine concentration 50 ng/mL. Related to Figure 5B.

Supplementary Table 11. List of genes from the IFNγ induction experiment, THP-1 macrophages, cytokine concentration 10 ng/mL. Related to Figure 5C.

Supplementary Table 12. List of genes from the IFNγ memory experiment, THP-1 monocytes, cytokine concentration 50 ng/mL. Related to Figure 6B.

Supplementary Table 13. List of genes from the IFNγ memory experiment, THP-1 with monocyte-to-macrophage transition, cytokine concentration 50 ng/mL. Related to Figure 6B.

Supplementary Table 14. List of genes from the LPS memory experiment, THP-1 with monocyte-to-macrophage transition, LPS concentration 100 ng/mL. Related to Figure 6B.

Supplementary Table 15. List of genes from the IFNγ priming experiment, THP-1 with monocyte-to-macrophage transition, cytokine concentration 50 ng/mL. Related to Figure 6E.

Supplementary Table 16. DNA oligonucleotides used in this work. Related to Materials and Methods.

Supplementary Table 17. DNA and protein sequences of STAT1 and STAT3 constructs for induction and purification from *Escherichia coli*. Related to Materials and Methods.

## Figure Legends

**Supplementary Figure 1. STAT3 recovers after SD36 treatment within 3 days after washout of the molecule. (A)** Expression level of STAT1 and STAT3 in naïve and IFNγ stimulated (50 ng/mL) HeLa cells. **(B)** Quantification (top) of GBP5 protein level (normalized to ACTB) from dot blot (bottom) showing the IFNγ memory in the HeLa cell line. **(C)** Western blot results (related to Figure 1D) showing STAT3 and STAT1 protein levels after removal of STAT3 with SD36 PROTAC (10µM), with or without IFNγ stimulation. ACTB – loading control. **(D)** Western blot results (related to Figure 1E) showing the dynamics of STAT3 removal and recovery in the HeLa cell line after treatment with the SD36 PROTAC (10µM). ACTB – loading control.

**Supplementary Figure 2. Reconstituted, unphosphorylated STAT3 exists in two conformations. (A)** Western blot results (related to Figure 2A) showing the effect low (1µM) and high (10µM) concentration of SD36 on STAT1 phosphorylation at positions S727 and Y701 as well as STAT1 and STAT3 total protein levels upon treatment with IFNγ (50 ng/mL) in THP-1 macrophages. ACTB – loading control. **(B)** Coomassie stained, SDS-PAGE results for STAT1 and STAT3 induction and purification from *Escherichia coli*. ctrl – without induction, NiNTA – after purification on a nickel column, GST cut – after on column removal of the GST-tag with Prescission Protease. **(C)** Results of a calibrated size exclusion measurement for STAT1 and STAT3 proteins expressed and purified from *Escherichia coli*. On top, table with theoretical and experimentally observed masses in kDa. At bottom, column plot with experimentally determined masses of proteins, as well as the standards used for column calibration. **(D)** STAT3 crosslinking results. Coomassie-stained SDS-PAGE gel (left) shows STAT3 crosslinked with 0.5µM (low) and 1µM (high) BS3. Bands analyzed by mass spectrometry are labeled. Mass spectrometry detected intra- and inter-molecular crosslinks (right).

**Supplementary Figure 3. Human primary monocytes respond to IFNγ in a concentration dependent manner. (A)** Flow cytometric analysis of HLA-DR, CD86, and CD80 expression on CD14-positive monocytes within PBMCs treated with increasing concentrations of IFNγ. **(B)** Comparison of HLA-DR expression in CD14-positve monocytes within PBMCs and for isolated CD14-positive monocytes treated with 2 ng/mL IFNγ, analyzed by flow cytometry. **(C)** Expression levels of genes identified in Figure 3B (GVINP1 and B7-H6) in male (top) and female (bottom) primary macrophages for different conditions of the IFNγ memory experiment (Figure 3A). **(D)** As in panel C, but for previously identified memory genes: GBP5 and GBP1.

**Supplementary Figure 4.1. Simplification of human monocyte-to-macrophage differentiation protocol. (A)** Measurement of efficacy of the simplified monocyte-to-macrophage differentiation protocol. (Top) Representative micrographs comparing the morphology of isolated monocytes and PBMCs after M-CSF-driven differentiation into macrophages. (Bottom) Flow cytometry analysis of monocyte (CD14) and macrophage (CD68) markers for isolated monocytes and PBMCs with or without M-CSF treatment. **(B)** Expression levels of IL32 and CCL7 genes (identified in Figure 4B) in male and female primary macrophages for different conditions of the IFNγ memory experiment (Figure 4A). **(C)** Expression levels of female specific memory gene CLEC5A in male and female primary macrophages for different conditions of the IFNγ memory experiment. **(D)** Expression levels of BATF2, TIFA, IFI44L and CXCL9 genes (identified in Figure 4C) in male and female primary macrophages for different conditions of the IFNγ memory experiment (Figure 4A).

**Supplementary Figure 4.2. Sex-bias in IFNγ memory.** Dot blot quantification of CXCL9 protein levels (normalized to ACTB) in primary macrophages from female (n = 9, top) and male (n = 10, bottom) donors across the IFNγ memory experiment (as in Figure 4A).

**Supplementary Figure 5. IFNγ memory is dependent on the cytokine concentration. (A)** Flow cytometry viability measurement of primary monocytes immediately after isolation (left), one day later (middle) and eight days into culture (right). **(B)** Expression levels of CXCL8, 1 & 3 genes (identified in Figure 5B) in THP-1 macrophages at the indicated concentrations of IFNγ (2, 10 & 50 ng/mL; see Figure 5A). **(C)** Quantification of CXCL9 protein level in THP-1 macrophages.

**Supplementary Figure 6. Monocyte-to-macrophage differentiation gates recall of IFNγ memory. (A)** Expression level of GBP5 obtained by RT-qPCR for different conditions of the IFNγ memory experiment in monocytes and during monocyte-to-macrophage transition in the memory window (as in Figure 6A). **(B)** Expression level of MMP8 and LOXL4 for different conditions of the IFNγ memory experiment in monocytes and during monocyte-to-macrophage transition in the memory window (as in Figure 6A).

