## Supplementary Figures 1-6 for "STAT3 dual role in sex-biased and differentiation-gated interferon gamma memory"

### SUPPLEMENTARY FIGURE 1

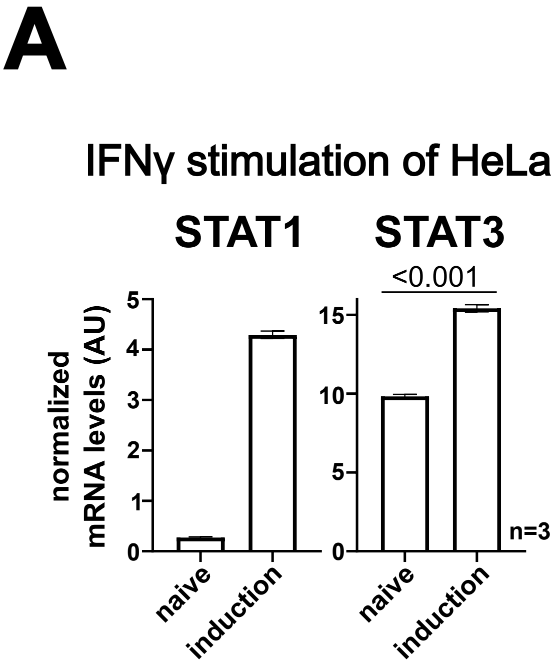


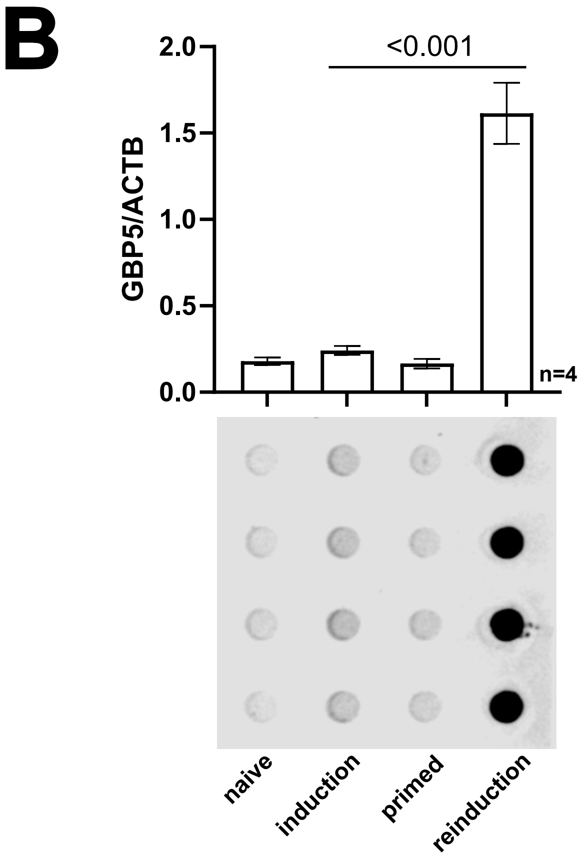


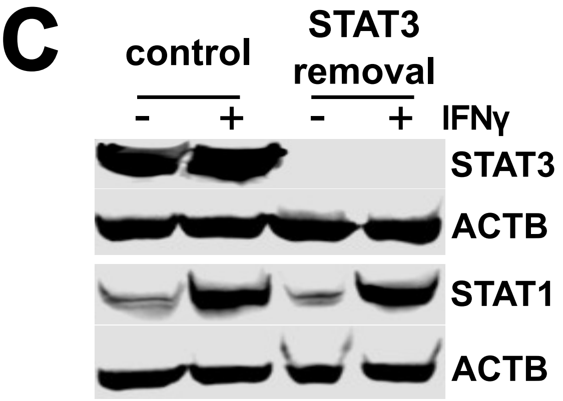


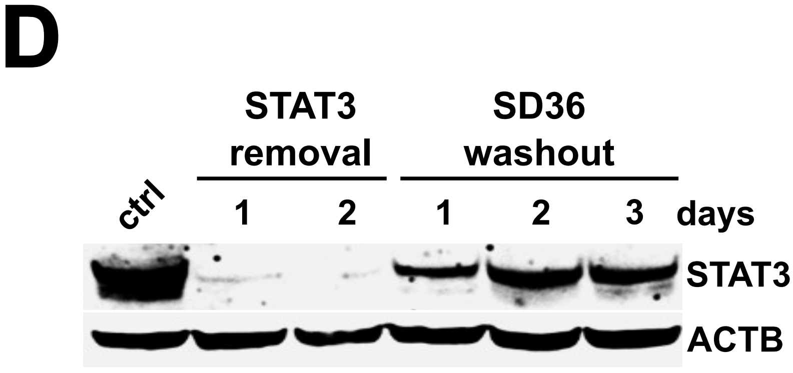

### SUPPLEMENTARY FIGURE 2

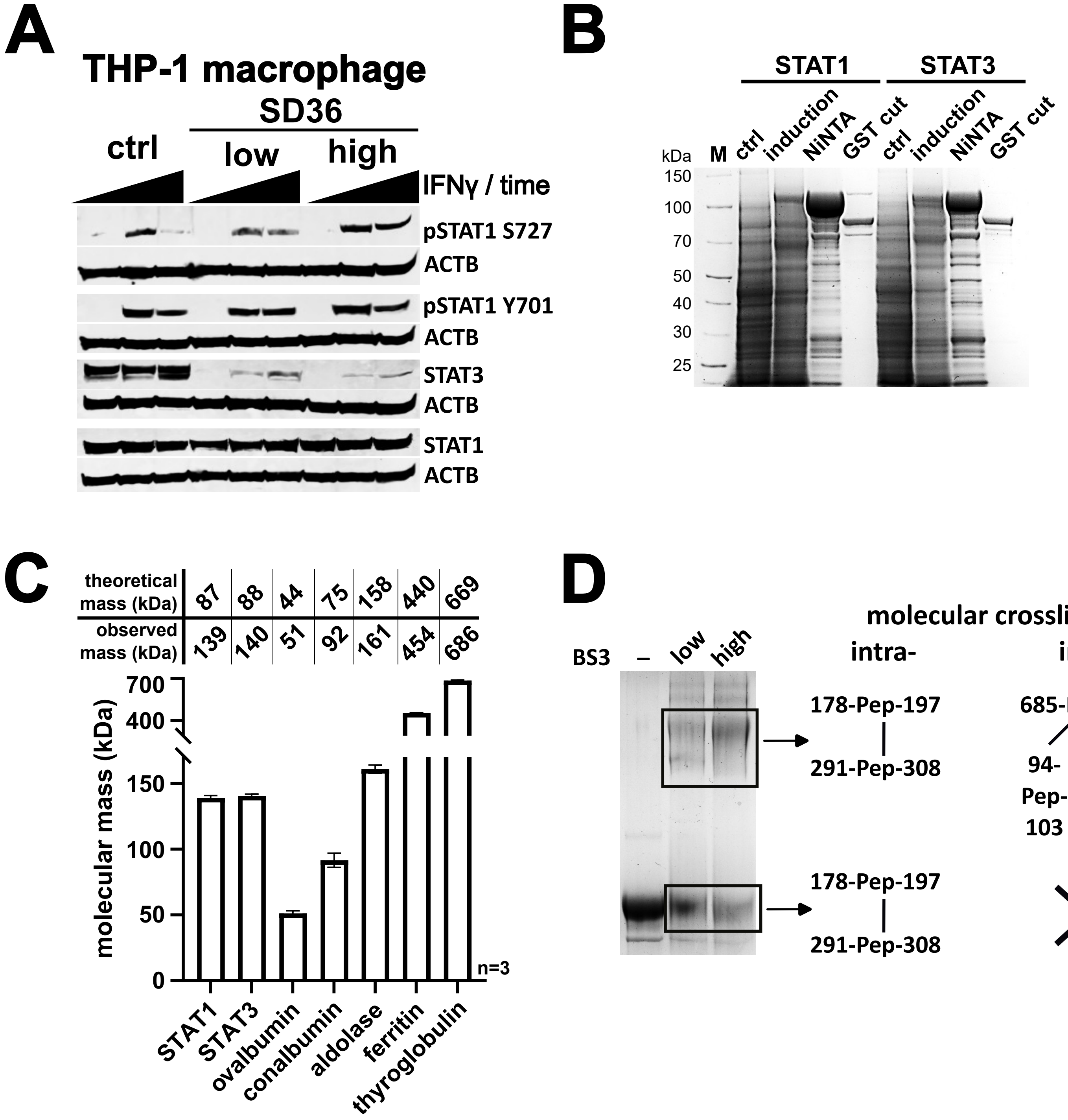

### SUPPLEMENTARY FIGURE 3

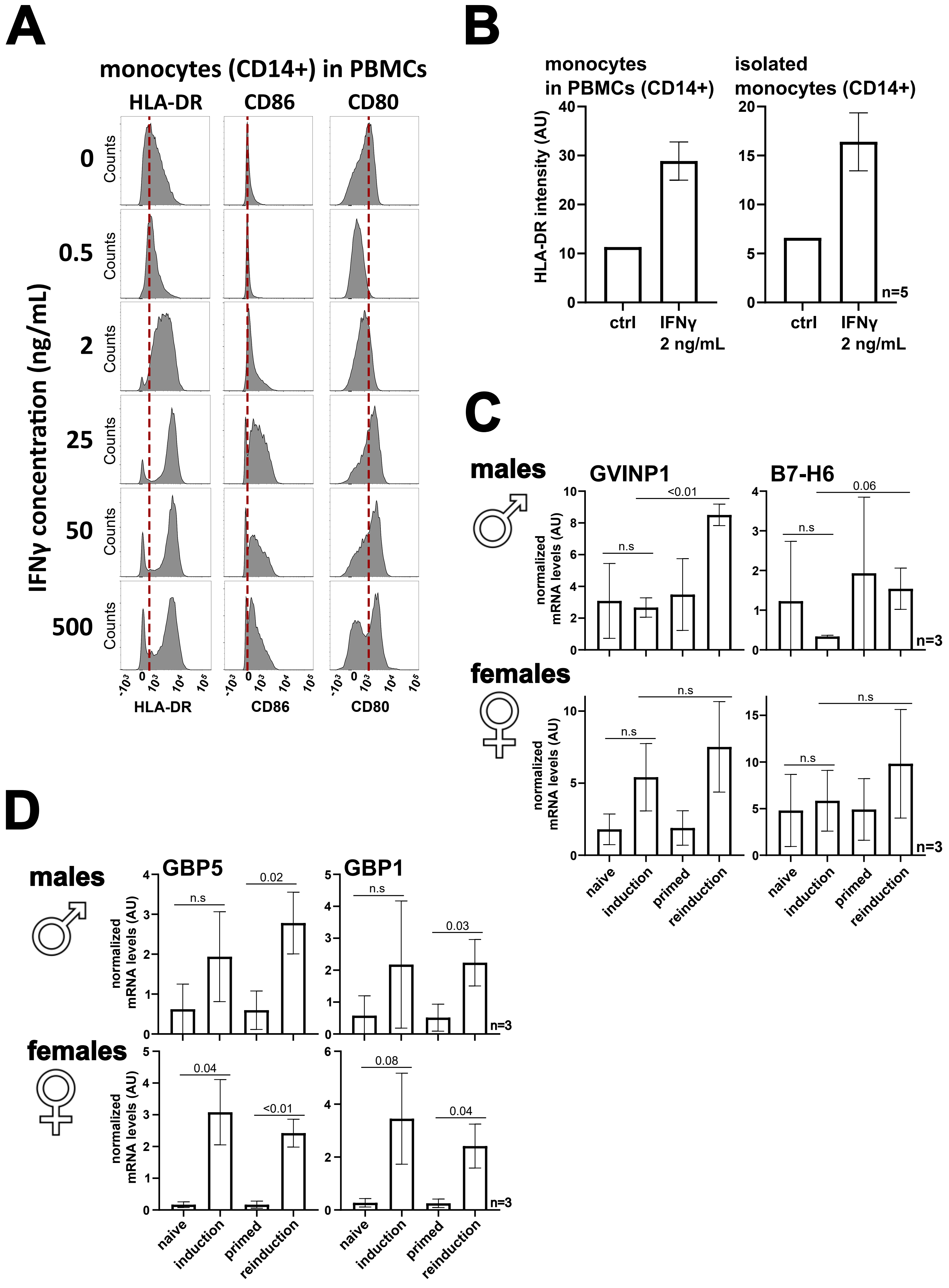

### SUPPLEMENTARY FIGURE 4.1

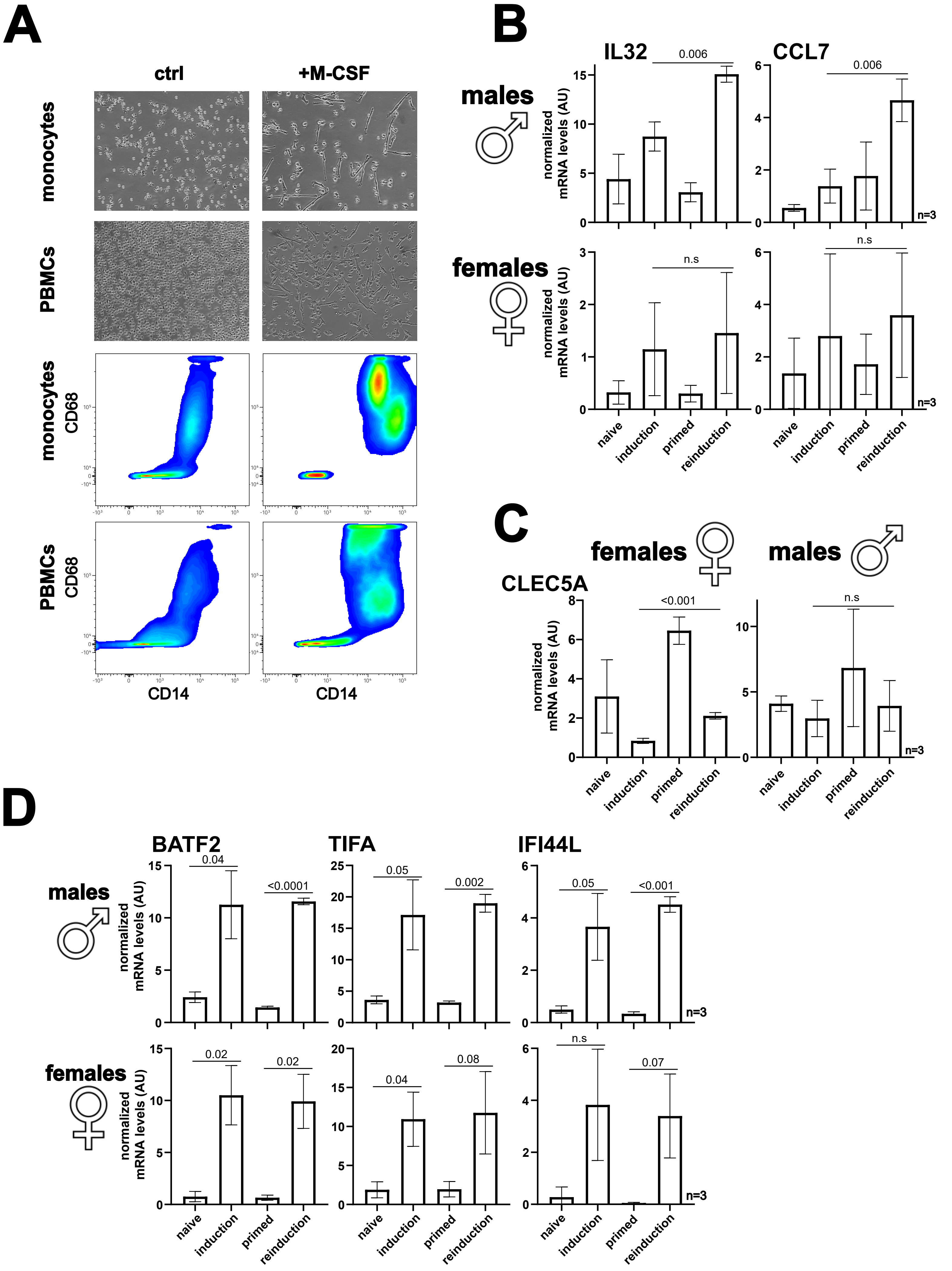

### SUPPLEMENTARY FIGURE 4.2

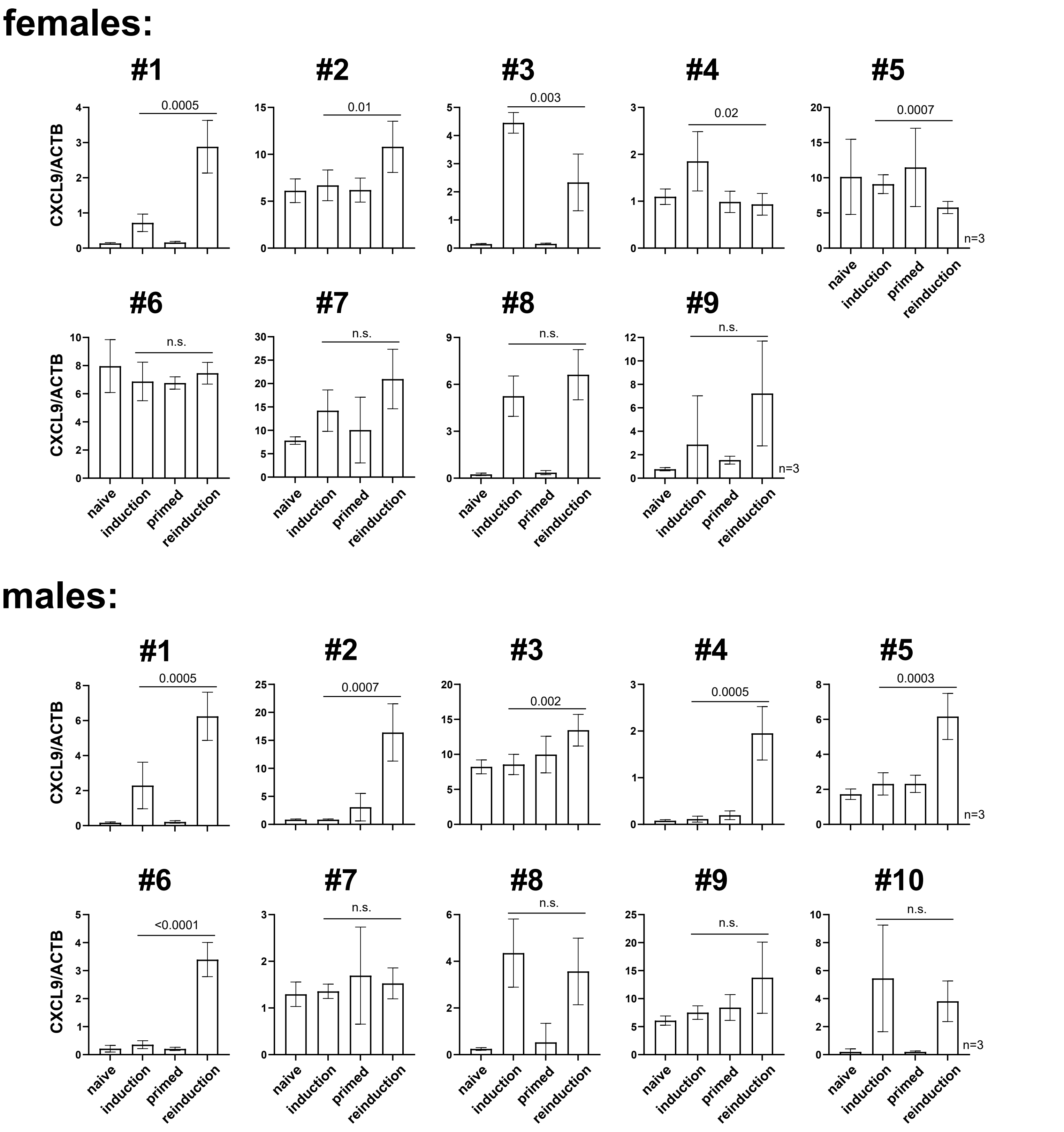

### SUPPLEMENTARY FIGURE 5

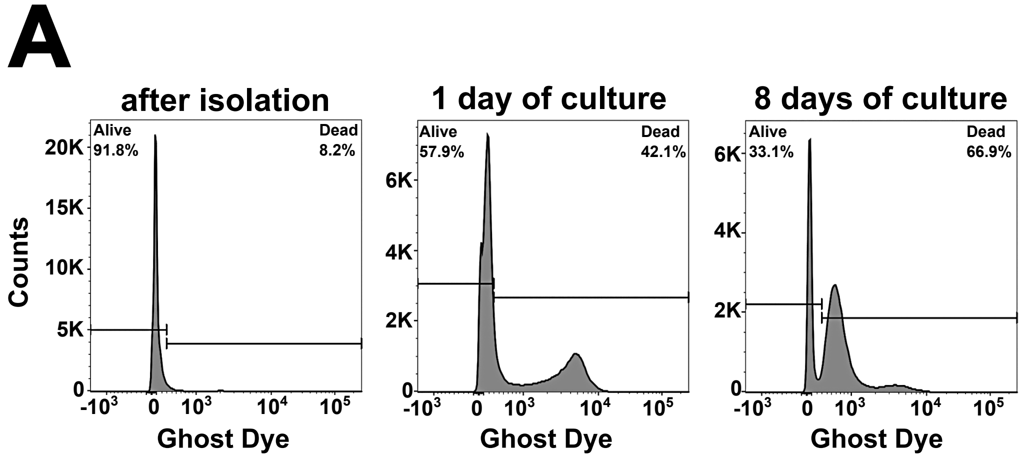


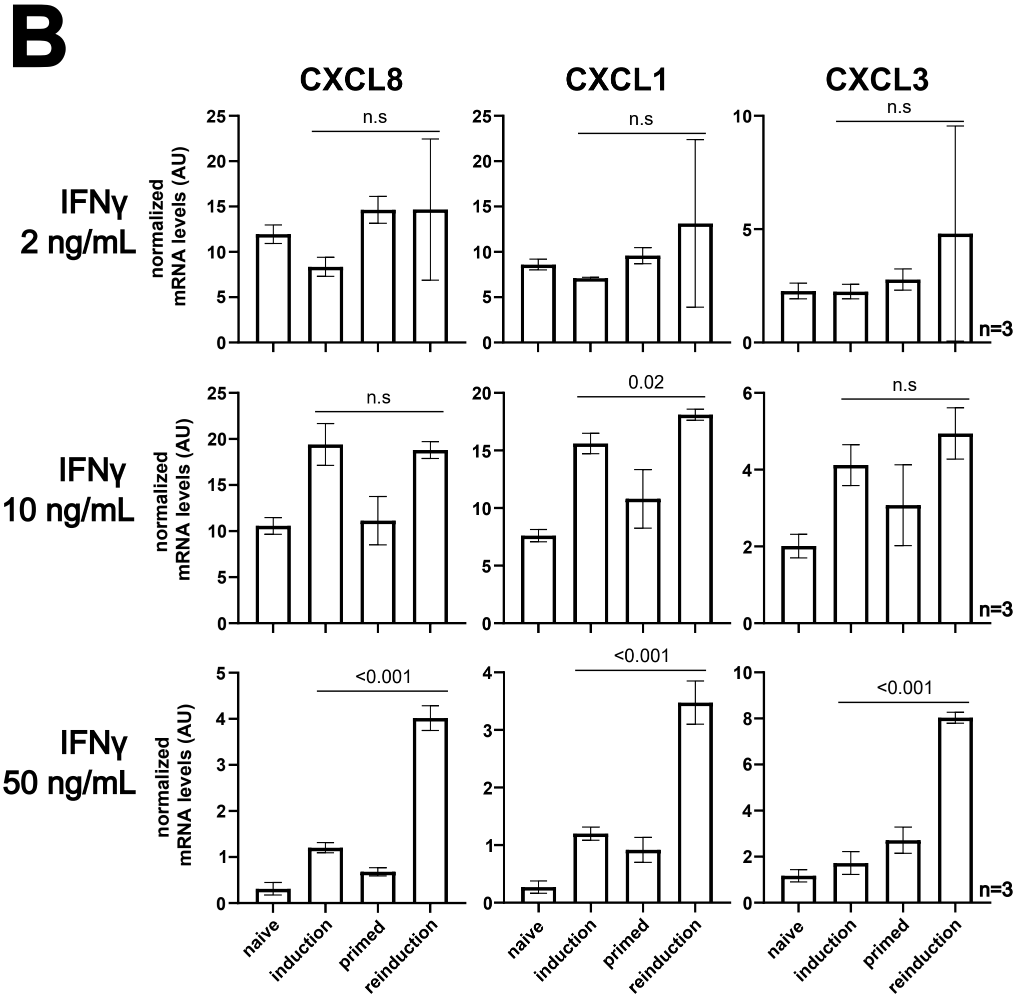


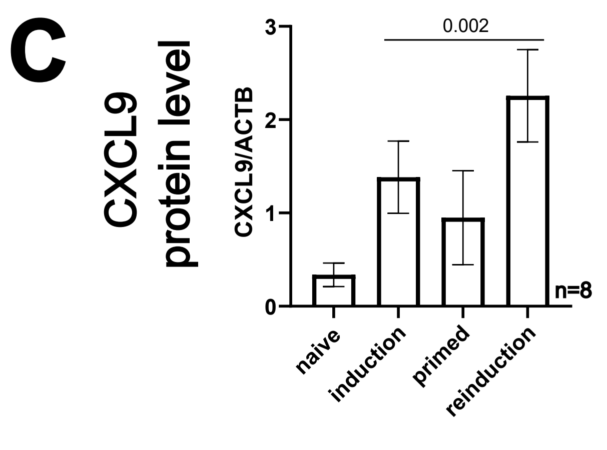

### SUPPLEMENTARY FIGURE 6

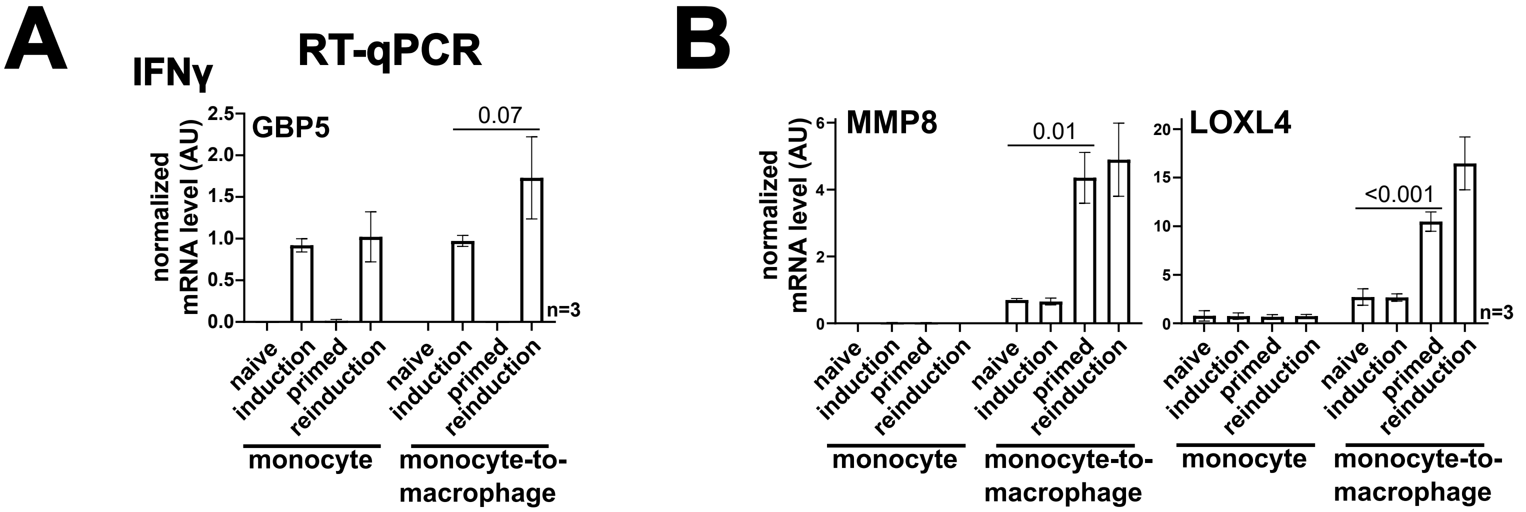
